# Mucosal vaccine adjuvant cyclic di-GMP differentiates lung moDCs into Bcl6^+^ and Bcl6^−^ mature moDCs to induce lung memory CD4^+^ T_H_ cells and lung T_FH_ cells respectively

**DOI:** 10.1101/2020.06.04.135244

**Authors:** Samira Mansouri, Divya S Katikaneni, Himanshu Gogoi, Lei Jin

## Abstract

Induction of lung T-cell responses, including memory CD4^+^ T_H_ and T_FH_ cells, are highly desirable for vaccines against respiratory infections. We recently showed that the non-migratory monocytes-derived DCs (moDCs) induced lung T_FH_ cells. However, the DCs subset inducing lung CD4^+^ memory T_H_ cells is unknown. Here, using conditional knockout mice and adoptive cell transfer, we first established that moDCs are essential for lung mucosal, but are dispensable for systemic, vaccine responses. Next, we showed that intranasal administration of adjuvant cyclic di-GMP differentiated lung moDCs into Bcl6^+^ and Bcl6^-^ moDCs promoting lung memory T_H_ cells and lung T_FH_ cells, respectively. Mechanistically, soluble TNF from lung TNFR2^+^ cDC2 subpopulation mediates the induction of lung Bcl6^+^ moDCs. Last, we designed fusion proteins targeting soluble or transmembrane TNF to lung moDCs and generated Bcl6^+^, Bcl6^-^ lung moDCs respectively. Together, our study revealed lung mature moDCs heterogeneity and showed a moDCs-targeting strategy to enhance lung mucosal vaccine responses.

## Introduction

Vaccination is the most cost-effective approach to fight infectious diseases. Approved vaccines mainly elicit antibody-mediated protection and do not generate strong mucosal responses. Vaccines that induce T-cell-mediated protection in lung mucosa would provide major health and economic benefits. Memory T cells include central memory T cells (T_CM_), effector memory T cells (T_EM_), and tissue-resident memory T cell (T_RM_). T_RM_ cells comprise a majority of memory T cells in the lung and play a crucial role in maintaining long-term protective immunity in lung mucosa(Mueller and Mackay, 2016; Sathaliyawala et al., 2013). Lung T_RM_ cells include CD4^+^ T_RM_ and CD8^+^ T_RM_ cells. While most studies focused on CD8^+^ T_RM_ cells, the induction of lung CD4^+^ T_RM_ cells is equally important.

Lung CD4^+^ T_RM_ cells are crucial for protection against influenza virus and *S.pneumoniae* infection(Smith et al., 2018; Teijaro et al., 2011). Swain’s group showed that lung memory CD4^+^ T cells protect against influenza through multiple synergizing mechanisms(McKinstry et al., 2012). Memory CD4^+^ T cells can accelerate primary CD8^+^ T cells responses(Krawczyk et al., 2007), activate innate immune cells to combat infections(Strutt et al., 2010), and secret T_H_ cytokines(McKinstry et al., 2010). The DCs subset promote the induction of lung CD4^+^ T_RM_ cells is unknown. Previous studies suggested that CD8^+^ DCs (cDC1) promote CD8^+^ T_RM_ cells(Iborra et al., 2016; Wakim et al., 2015). Several recent studies showed that monocytes promote the generation of CD8^+^ T_RM_ in the lung(Desai et al., 2018; Dunbar et al., 2019; Thompson et al., 2019). These monocytes do not express MHC class II(Dunbar et al., 2019). Thus, they are unlikely to induce lung CD4^+^ T_RM_ cells.

CD4^+^ T_RM_ and CD8^+^ T_RM_ cells share a core transcription signature such as CD69, CCR6 and CD49a(Kumar et al., 2017; Schreiner and King, 2018; Wilk and Mills, 2018). Nevertheless, key differences exist between CD4^+^ vs. CD8^+^ T_RM_. First, CD4^+^ T_RM_ far outnumbers CD8^+^ T cells in the lung(Sathaliyawala et al., 2013; Yang et al., 2011). Second, CD4^+^ T_RM_ cells are more polyclonal than CD8^+^ T_RM_ cells(Kumar et al., 2017; Lees and Farber, 2010; Stockinger et al., 2006). Third, CD8^+^ T_RM_ cells typically localize in the epithelial layers of barrier tissues and do not form clusters or lymphoid-like structures(Takamura et al., 2016). In contrast, CD4^+^ T_RM_ cells typically localize in cell clusters or ectopic lymphoid structures(Hondowicz et al., 2016; Turner et al., 2014). In the lung, most CD4^+^ T_RM_ cells are in B cell follicles and, by T cell areas(Hondowicz et al., 2016; Shinoda et al., 2012; Takamura et al., 2016; Turner et al., 2014), a lymphoid-like structure called inducible bronchus-associated lymphoid tissues (iBALT). Thus, lung CD4^+^ T_RM_ cells likely have a different induction mechanism from the lung CD8^+^ T_RM_ cells.

Lung T_RM_ cells (both CD4^+^ and CD8^+^ T_RM_) require local antigen presentation(Bautista et al., 2016; Haddadi et al., 2019; McKinstry et al., 2014; McMaster et al., 2018). In contrast, T_RM_ cells in the intestine, genital tract, or skin, do not require the tissue antigen recognition(Casey et al., 2012; Mackay et al., 2012). Swain’s group first reported that CD4^+^ T cell memory formation depends on re-engagement of CD4^+^ T cells with APCs at their effector stage (“Signal 4”), day 5-8 of their response(Bautista et al., 2016; McKinstry et al., 2014). Lung CD8^+^ T_RM_ establishment also requires cognate antigen recognition in the lung(McMaster et al., 2018). It is unclear which lung APCs provide the “Signal 4” for lung CD4^+^ T_RM_ induction at this stage of vaccination or infection.

Monocyte-derived dendritic cells (moDCs) are a DC subset that rapidly accumulates at the site of inflammation(Randolph et al., 1998). As a result, moDCs are found virtually in any diseases with substantial inflammation(Chow et al., 2017). Though previously known as TNF/iNOS-Producing Dendritic Cells (Tip-DCs)(Serbina et al., 2003) for their roles in promoting inflammation, these monocyte-derived Tip-DCs are likely not DCs because they were not essential for T cell priming(Serbina et al., 2003). Bona fide moDCs are highly capable of antigen uptake, processing, presentation, and priming CD4^+^ and CD8^+^ T cells(Chow et al., 2017). For example, tumor-infiltrating moDCs prime CD8^+^ T cells and induce anti-tumor immunity(Sharma et al., 2018). CCR2^+^ moDCs are critical for T_H_17 induction and the development of experimental autoimmune encephalomyelitis(Ko et al., 2014). During cutaneous *L. major* infection, moDCs could induce T_H_1 polarization(Leon et al., 2007). moDCs were also sufficient to induce T_H_2 immunity by high-dose HDM(Plantinga et al., 2013). Last, we recently showed that moDCs induce T_FH_ cells in the lung by mucosal adjuvant cyclic di-GMP(Mansouri et al., 2019). Thus, there is a significant degree of plasticity and heterogeneity in moDCs *in vivo*. The underlying mechanism for moDCs plasticity and heterogeneity is unknown(Chow et al., 2017).

In this report, we study the role of lung moDCs in generating lung mucosal CD4^+^ memory T_H_ cells, including lung CD4^+^ T_RMs_. We found that moDCs are specifically required for the generation of CD4^+^ memory T_H_ cells in lung mucosa but not in the systemic compartments. Furthermore, we identified new differentiated moDCs subpopulations responsible for CD4^+^ memory T_H_ cells and T_FH_ induction in lung mucosa, respectively.

## Results

### CCR2^-/-^ mice selectively lose cyclic di-GMP adjuvanticity in lung mucosa

Cyclic di-GMP (CDG) is a mucosal adjuvant eliciting balanced protective immunity in the systemic and mucosal compartments(Allen et al., 2018; Blaauboer et al., 2014; Blaauboer et al., 2015; Ebensen et al., 2007; Mansouri et al., 2019). We previously showed that moDCs promote lung mucosal T_FH_ and IgA responses(Mansouri et al., 2019). To further evaluate the role of moDCs on CDG adjuvanticity, we used CCR2^-/-^ mice, which have decreased numbers of lung moDCs (Figure S1A-S1B). We immunized (*i.n*.) CCR2^-/-^ mice with CDG/PspA and examined systemic, lung mucosal adjuvant responses on day 28. We found that CCR2^-/-^ mice had unaltered anti-PspA serum IgG but lacked anti-PspA IgA in the BALF (Figure S1C). We then examined memory T_H_1/2/17 responses in the spleen and lung from immunized CCR2^-/-^ mice by *ex vivo* recall. Again, CCR2^-/-^ mice retained memory T_H_ responses in the spleen but not in the lung (Figure S1D-S1E). Thus, CDG induced lung mucosa adjuvant responses could be uncoupled from systemic responses. moDCs may be specifically needed for promoting adjuvant responses in lung mucosa.

### moDCs are not activated in RelA^fl/fl^CD11c^cre^ mice, which had selective loss of CDG adjuvant-induced lung mucosal IgA and lung memory CD4 T cells responses

We previously showed that CDG activates RelA of the NF-κB signaling in moDCs(Mansouri et al., 2019). We thus generated RelA^fl/fl^CD11c^cre^ mice to examine the role of moDCs activation in CDG adjuvanticity in lung mucosa. Unlike the CCR2^-/-^ mice, RelA^fl/fl^CD11c^cre^ mice had normal numbers of moDCs (Figure S2A). However, RelA^fl/fl^CD11c^cre^ mice had no antigen-specific IgA in BALF while maintained antigen-specific IgG production in the serum (Figure 1A-1B). The RelA^fl/fl^CD11C^cre^ mice also did not generate memory T_H_ responses in the lung while retained memory T_H_ responses in the spleen (Figure 1C-1D). RelA^fl/fl^CD11c^cre^ mice had a selective defect on CDG-induced moDCs activation (Figure 1E, S2B). Consistently, CDG induced T_FH_ cells, CD4^+^ T_RM_ cells, and GC B cells were reduced in the lung (Figure S2C-2F).

**Figure 1.**
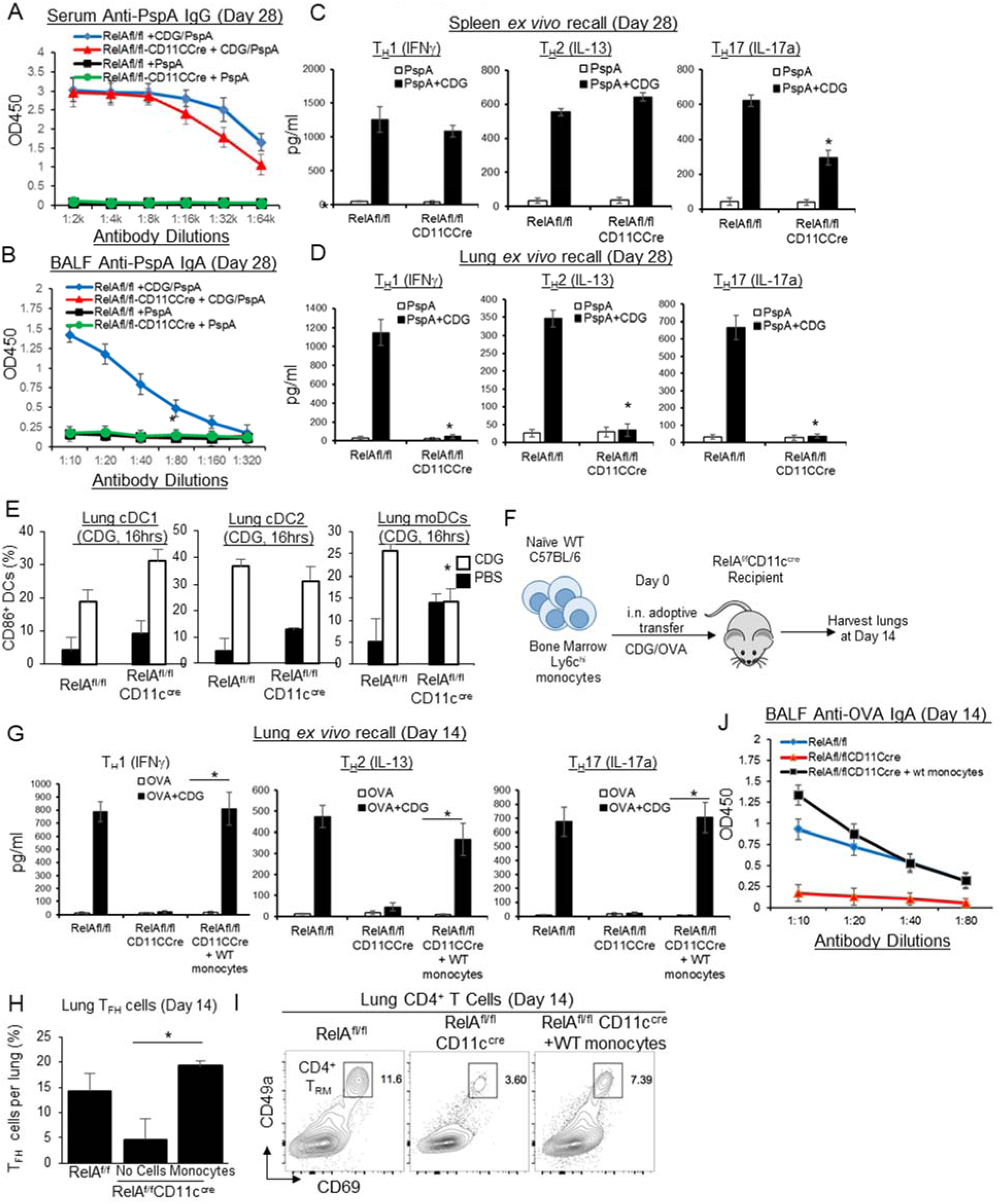
moDCs promote CDG adjuvant-induced lung mucosal IgA and CD4 memory T_H_ responses. **A-B**. RelA^fl/fl^ and RelA^fl/fl^CD11c^cre^ mice were immunized (*i.n*.) with two doses (14 days apart) of PspA or PspA plus CDG (5μg). Anti-PspA IgG in serum (**A**) and IgA in BALF (**B**) were determined by ELISA 28 days post-immunization. (n=3mice/group) Data are representative of three independent experiments. **C-D**. Lung cells (**D**) or splenocytes (**C**) from immunized RelA^fl/fl^ and RelA^fl/fl^CD11c^cre^ mice (**A**) were recalled with 5μg/ml PspA for 4 days in culture. Cytokines were measured in the supernatant by ELISA. **E**. Frequency of activated (CD86^+^) pulmonary DC subsets in RelA^fl/fl^ and RelA^fl/fl^CD11c^cre^ mice administered (*i.n*.) with PBS or 5µg CDG for 16 hours. (n=3mice/group). Data are representative of two independent experiments. **F**. Experimental design for adoptive transfer. RelA^fl/fl^CD11c^cre^ mice (n=3mice/group) were immunized with CDG/OVA. WT bone marrow Ly6C^hi^ monocytes (1.5 million total cells) were transferred (*i.n*.) at 30mins, 2hrs and 4hrs post-immunization. Mice were harvested on day 14. **G**. Lung cells from (**F**) were recalled with 2μg/ml OVA for 4 days in culture. Cytokines were measured in the supernatant by ELISA. Data are representative of three independent experiments. **H**. Frequency of lung T_FH_ from (**F**). Data are representative of three independent experiments. **I**. CD4^+^CD69^+^CD49a^+^ T_RM_ were determine by flow cytometry from (**F**). Data are representative of three independent experiments. **J**. BALF anti-OVA IgA on day 14 from (**F**) were determined by ELISA. Data are representative of three independent experiments. Graphs represent the mean with error bars indication S.E.M. *P* values determined by one-way ANOVA Tukey’s multiple comparison test (**C, D, H**) or unpaired student *t*-test (**E, G**). \**P*<0.05

### moDCs mediate CDG adjuvant-induced lung IgA and CD4+ memory T cells responses

RelA^fl/fl^CD11c^cre^ mice deleted RelA in DCs and alveolar macrophages. We thus generated RelA^fl/fl^LysM^cre^ mice to delete RelA in myeloid cells, including macrophage, monocytes, and monocyte-derived cells, *e.g*., moDCs. Similar to RelA^fl/fl^CD11c^cre^ mice, RelA^fl/fl^LysM^cre^ mice lost IgA and memory T_H_ cells in lung mucosa but not in the systemic compartments (Figure S3A-S3C). Together, the data suggested that RelA in moDCs, not cDCs, mediates CDG adjuvant responses in lung mucosa.

moDCs are differentiated from Ly6C^hi^ monocytes(Randolph et al., 1998). We isolated bone marrow Ly6C^hi^ monocytes from C57BL/6J mice and adoptively transferred (*i.n*.) them into RelA^fl/fl^CD11c^cre^ mice immunized with CDG/OVA (Figure 1F). The procedure was repeated once after 14 days. We examined lung mucosal immune responses 14 days after the last immunization. Indeed, RelA^fl/fl^CD11C^cre^ mice received WT monocytes had restored lung CD4^+^ T_RM,_ T_FH_ cells, and lung memory T_H_ responses (Figure 1G-1I). Adoptively transferred WT monocytes also restored OVA-specific IgA in BALF (Figure 1J). Together, the data indicated that lung moDCs mediate the generation of lung mucosa-specific IgA and lung CD4^+^ memory T_H_ cells.

### moDCs differentiate into Bcl6^+^ moDCs in the lung

How do moDCs simultaneously promote lung IgA/T_FH_ and lung memory CD4^+^ T_H_ responses? We hypothesized that moDCs might differentiate into functionally distinct mature lung moDCs to induce T_FH_ and CD4 memory T_H_ cells respectively in the lung.

We first noticed that on day 14 post-immunization, lungs, not lung draining lymph nodes, contain a population of moDCs with elevated Bcl6 (transcriptional repressor B-cell CLL/lymphoma 6) expression (Figure 2A). Bcl6 is the master transcriptional factor for T_FH_ and GC B cells. A population of lung cDC2 also express Bcl6 (Figure 2A). The Bcl6^+^ cDC2 population is mainly TNFR2^-^ cDC2. (Figure 2A, S4A). Lung cDC1 does not express Bcl6 on day 14 post-immunization (Figure 2A-2B). Bcl6 expression is lower in lung Bcl6^+^ moDCs than in lung T_FH_ or GC B cells (Figure S4B).

**Figure 2.**
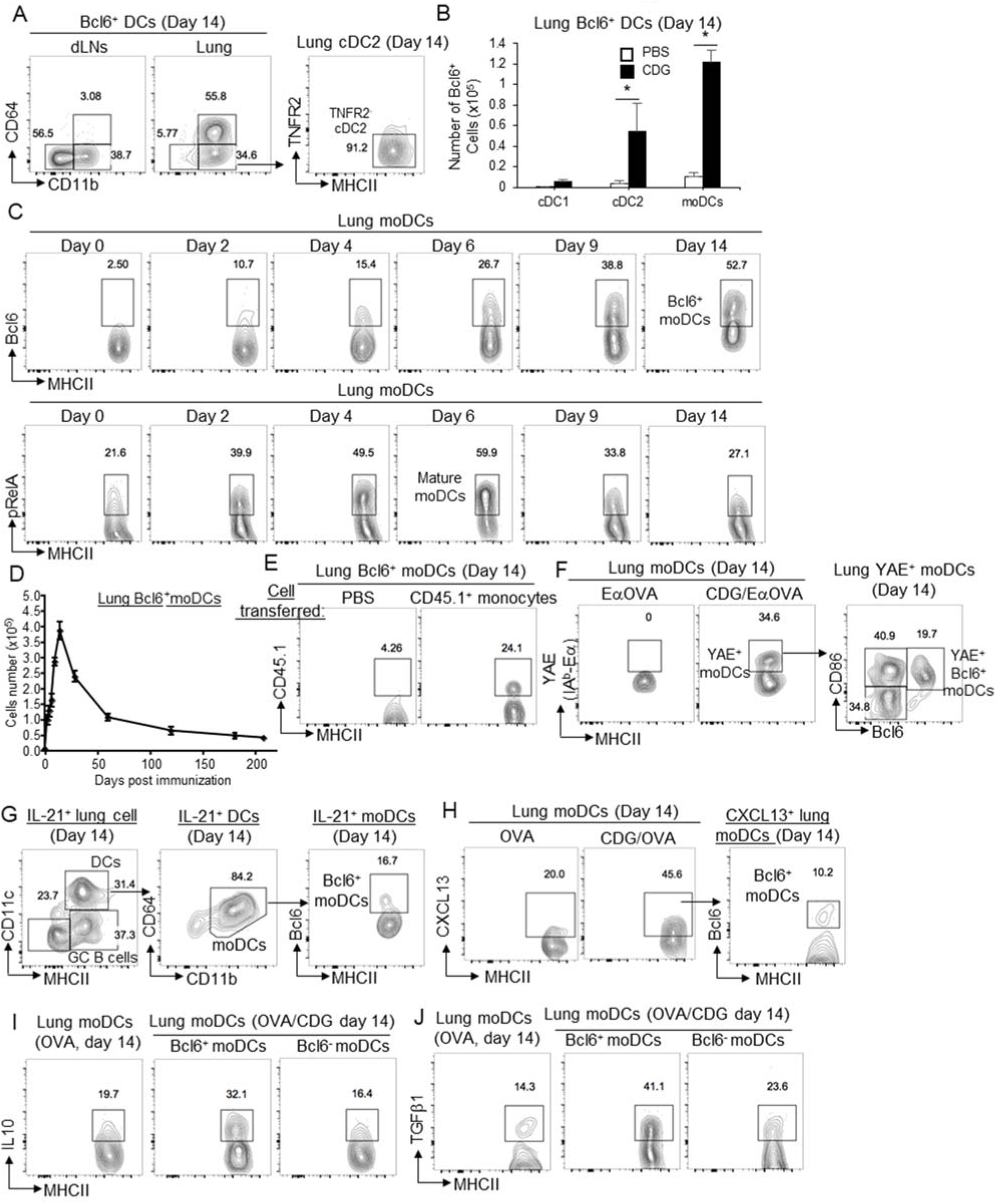
moDCs differentiate into Bcl6^+^ and Bcl6^-^ mature moDCs in the lung mucosa. **A**. Flow cytometry analysis of Bcl6 expressing DCs in the mediastinal lymph nodes and lungs of WT mice immunized with CDG/OVA. Lungs were harvested on day 14. (n=3mice/group) Data are representative of two independent experiments. **B**. The absolute numbers of Bcl6^+^ pulmonary DC subsets in the lungs of WT mice immunized with CDG/OVA on day 14. (n=3mice/group) Data are representative of two independent experiments. **C**. Flow cytometry analysis of Bcl6 (top) and pRelA (bottom) expression in lung moDCs of WT mice immunized with CDG/OVA. Lungs were harvested at different time points. (n=3mice/group) Data are representative of two independent experiments. **D**. The kinetics of the absolute number of Bcl6^+^ moDCs in WT mice at different time points following immunization (*i.n*.) with CDG/OVA. (n=3mice/group) Data are representative of two independent experiments. **E**. WT mice receiving CD45.1^+^ monocytes (*i.n*.) were immunized with CDG/NP_6_CGG. On day 14, CD45.1^+^Bcl6^+^ moDCs were determined by Flow cytometry. (n=3mice/group) Data are representative of two independent experiments. **F**. WT mice were immunized with Eα-OVA or Eα-OVA/CDG (5µg). YAE^+^ moDCs were determined by flow cytometry in the lung on day 14. (n=3mice/group) Data are representative of three independent experiments. **G**. Analysis of IL-21^+^ cells in the lungs of IL-21-VFP reporter mice on day 14 post-immunization with CDG/OVA (*i.n*.). (n=3mice/group) Data are representative of two independent experiments. **H**. Flow cytometry analysis of CXCL13^+^ lung moDCs in C57BL/6J mice on day 14 post-immunization (*i.n*.) with CDG/OVA. (n=3mice/group) Data are representative of two independent experiments. **I-J**. Flow cytometry analysis of IL-10 (**I**), and TGFβ1 (**J**) production by lung moDCs in C57BL/6J mice on day 14 post-CDG/OVA immunization. (n=3mice/group) Data are representative of two independent experiments. Graphs represent the mean with error bars indication s.e.m. *P* values determined by unpaired student *t*-test. \**P*<0.05, \*\**P*<0.001, \*\*\**P*<0.0001.

Next, we examined the kinetics of moDCs differentiation in the lung during CDG immunization. The activation of moDCs, indicated by pRelA and CD80 peaked on day 6 post-immunization (Figure 2C, S4D) before the appearance of lung T_FH_ and GC B cells (day 9) (Figure S4E-S4F). Intriguingly, Bcl6^+^ moDCs differentiation had different kinetics from moDCs activation. Bcl6^+^ moDCs appeared early on day 2 and kept increasing to day 14, even when moDCs activation was waning (Figure 2C). We found that, on day 14 post-immunization, ∼52% of lung moDCs expressed Bcl6 (Figure 2C). Unexpectedly, the Bcl6^+^ moDCs were long-lasting, detected >200 days post-immunization (Figure 2D).

To confirm that the Bcl6^+^ moDCs were indeed monocyte-derived, we adoptively transferred (*i.n*.) CD45.1^+^ WT Ly6C^hi^ monocyte into a C57BL/6J mouse immunized with CDG. On day 14 post-immunization, we identified CD45.1^+^ Bcl6^+^ moDCs (Figure 2E). Thus, these Bcl6^+^ moDCs are indeed monocytes-derived. Alveolar macrophages can be monocytes-derived and are CD11C^+^MHC II^+^. We, thus, examined alveolar macrophages from BALF in immunized mice and found that alveolar macrophages in immunized mice were Bcl6^-^ (Figure S4C). Thus, the Bcl6^+^ monocyte-derived cells were not BAL AM.

We reasoned that lung Bcl6^+^ moDCs could be induced by other stimuli. Indeed, adjuvants chitosan and cholera toxin (CT) induce lung Bcl6^+^ moDCs (Figure S5A). In comparison, adjuvant CpG did not induce lung Bcl6^+^ moDCs (Figure S5A). A previous study found that HDM treatment in mice generated CD4^+^ memory T_H_ cells in the lung(Turner et al., 2018). We found that HDM-treatment induced Bcl6^+^ moDCs in the lung (Figure S5B). Last, as expected, CDG-induced lung Bcl6^+^ moDCs were significantly reduced in RelA^fl/fl^CD11c^cre^ mice (Figure S5C).

### Both Bcl6^+^ and Bcl6^-^ lung moDCs present antigen, express co-stimulators but produce different cytokines on day 14 post-immunization

We then asked if the lung Bcl6^+^ DCs presented antigen at the effector stage (day 14) *in vivo*. We immunized C57BL/6J mice with CDG plus a fusion protein of Eα peptide (aa52-68) with ovalbumin (Eα-OVA). On day 14 post-immunization, we detected MHC class II-Eα peptide-specific DCs with the YAE mAb (Figure 2F). The YAE^+^ moDCs contained both Bcl6^+^ and Bcl6^-^ populations that were CD86^+^ (Figure 2F). Thus, both Bcl6^+^ and Bcl6^-^ moDCs were mature APCs on day 14 post-immunization.

We next asked if the Bcl6^+^ and Bcl6^-^ moDCs produce cytokines, the “Signal 3”. IL-21 is a major T_FH_-inducing cytokine. Using an IL-21 reporter mouse, we found that lung moDCs were the main IL-21-producing lung DCs in the lung on day 14 post-immunization (Figure 2G). However, the IL-21^+^ moDCs were mainly Bcl6^-^ moDCs (Figure 2G). CXCL13 is essential for GC formation. We found that on day 14 post-immunization, moDCs produced CXCL13 (Figure 2H). However, most CXCL13^+^ moDCs were Bcl6^-^ moDCs (Figure 2H). In contrast, the lung Bcl6^+^ moDCs produce IL-10, TGFβ1 (Figure 2I-2J) cytokines that are critical for the induction of T_RM_s(Nath et al., 2019; Thompson et al., 2019). The lung Bcl6^+^ moDCs also produce T_H_ promoting cytokines IL-23 and IL-4 (Figure S6). Together, the data indicated that lung moDCs differentiated into mature Bcl6^+^ and Bcl6^-^ moDCs on day 14 post-immunization and may promote different T-cells responses in the lung.

### Bcl6^fl/fl^LysM^cre^ mice are defective in lung moDCs development and lack CDG-induced lung mucosal IgA and lung CD4^+^ memory T_H_ responses

To understand the functional significance of differentiated moDCs populations *in vivo*, we generated Bcl6^fl/fl^LysM^cre^ and Bcl6^fl/fl^CD11c^cre^ mice. Bcl6^fl/fl^LysM^cre^ mice delete Bcl6 in myeloid cells. The cDCs population in the Bcl6^fl/fl^LysM^cre^ mice were unaltered (Figure S7A-S7B). The Bcl6^fl/fl^LysM^cre^ mice had expanded lung neutrophils and Ly6C^hi^ monocytes populations (Figure S7C) as previously reported(Zhu et al., 2019). However, Bcl6^fl/fl^LysM^cre^ mice had reduced lung moDCs (Figure S7A-S7B) similar to the CCR2^-/-^ mice (Figure S1B). Consequently, the Bcl6^fl/fl^LysM^cre^ mice lost lung mucosal-specific IgA production (Figure S7E) and lung memory T_H_ responses (Figure S7G). Spleen memory T_H_ responses and serum antigen-specific IgG in Bcl6^fl/fl^LysM^cre^ mice were comparable to the Bcl6^fl/fl^ mice 4 months post-CDG immunization (Figure S7D, S7F). Last, CDG immunized Bcl6^fl/fl^LysM^cre^ mice did not have lung T_FH_ and lung T_RM_ cells (Figure S7H, S7I). Thus, Bcl6 expression in the myeloid cells is crucial for Ly6C^hi^ monocytes to differentiate into moDCs in the lung. Like the CCR2^-/-^ or RelA^fl/fl^CD11c^cre^ mice, Bcl6^fl/fl^LysM^cre^ mice lack functional lung moDCs and can not promote lung mucosal memory T_H_ or T_FH_ cells.

### Bcl6^fl/fl^CD11c^cre^ mice lack lung cDC1 but have functional cDC2 to mediate CDG adjuvanticity in the systemic compartment

Bcl6 is a pleiotropic transcriptional factor. To bypass the influence of Bcl6 on lung moDCs development, we generated Bcl6^fl/fl^CD11c^cre^ mice to ablate Bcl6 after CD11c expression and hoped lung moDCs would be developed.

The Bcl6^fl/fl^CD11c^cre^ mice will delete the Bcl6 gene in CD11c^+^ cells, including cDCs and AM. A previous study found that BCL6 protein expression is elevated in peripheral pre-cDCs, especially cDC1(Zhang et al., 2014). We observed that Bcl6^fl/fl^CD11C^cre^ mice lack the lung cDC1 population (Figure S8A), suggesting that Bcl6 is required for lung cDC1 development. The lung cDC2 also decreased in Bcl6^fl/fl^CD11C^cre^ mice though was not statistically significant (Figure S8B). Lung cDC2 consists of functionally distinct TNFR2^+^ and TNFR2^-^ cDC2 populations (Figure S4A)(Mansouri et al., 2019). Notably, the TNFR2^-^ cDC2 population was significantly reduced in the Bcl6^fl/fl^CD11c^cre^ mice (Figure S8B). It is worth noting that the TNFR2^-^ cDC2, not the TNFR2^+^ cDC2, express Bcl6 on day 14 post-CDG immunization (Figure 2A). The significance of the lung Bcl6^+^ TNFR2^-^ cDC2 is unknown.

cDC2 is essential for CDG mucosal adjuvanticity while cDC1 is dispensable(Mansouri et al., 2019). We next examined the function of lung cDC2 in the Bcl6^fl/fl^CD11c^cre^ mice. Mice were treated (*i.n*.) with CDG for 16hrs. Lung cDC2 activation was determined by CD86 and CCR7 expression. The CD86 and CCR7 on CDG-activated lung cDC2 were comparable between Bcl6^fl/fl^ and Bcl6^fl/fl^CD11c^cre^ mice (Figure S8C-S8D). However, cDC2 from the Bcl6^fl/fl^CD11c^cre^ mice had elevated basal CD86 and CCR7 expression (Figure S8C-S8D). cDC2 is essential for CDG adjuvanticity(Mansouri et al., 2019). To further determine the function of cDC2 in Bcl6^fl/fl^CD11c^cre^ mice, we measured CDG adjuvanticity in the systemic compartments. Four months after CDG immunization, the Bcl6^fl/fl^CD11c^cre^ mice had unaltered antigen-specific serum IgG (Figure S8E) and memory T_H_ responses in the spleen (Figure S8F). Thus, the lung cDC2 in the Bcl6^fl/fl^CD11c^cre^ mice can promote CDG adjuvanticity and are functional.

### CDG adjuvant induces lung T_FH_ cells, lung IgA responses but not lung memory T_H_ responses in Bcl6^fl/fl^CD11c^cre^ mice

We then examined moDCs and lung mucosal responses in the Bcl6^fl/fl^CD11c^cre^ mice. Unlike the Bcl6^fl/fl^LysM^cre^ mice, Bcl6^fl/fl^CD11C^cre^ mice had increased numbers of lung moDCs at the steady-state (Figure S8B). These moDCs also had elevated CD86 expression (Figure S8G). Notably, lung moDCs from Bcl6^fl/fl^CD11c^cre^ mice did not upregulate Bcl6 expression on day 14 post-CDG immunization (Figure S8H). Unexpectedly, the basal Bcl6 expression in lung moDCs was elevated in the Bcl6^fl/fl^CD11c^cre^ mice compared to the Bcl6^fl/fl^ mice (Figure S8I). This elevated basal Bcl6 expression in lung moDCs was consistently observed in all Bcl6^fl/fl^ CD11c^cre^ mice we examined (>20 mice). Bcl6 is required for moDCs development (Figure S7A). We speculated that lung moDCs developed in the Bcl6^fl/fl^CD11c^cre^ mice by escaping the Bcl6 deletion and enhanced basal Bcl6 expression. Nevertheless, CDG immunization did not induce Bcl6 expression in lung moDCs from Bcl6^fl/fl^CD11c^cre^ mice (Figure S8H).

We next examined moDCs function in the Bcl6^fl/fl^CD11c^cre^ mice. moDCs are essential for promoting lung mucosal CDG responses. We immunized (*i.n*.) Bcl6^fl/fl^CD11c^cre^ mice with CDG/OVA twice at the two-weeks interval and examined mucosal lung vaccine adjuvant responses. Unlike the Bcl6^fl/fl^LysM^cre^ mice (Figure S7), the Bcl6^fl/fl^CD11c^cre^ mice had unaltered lung T_FH_ cells, GC B cells, or IgA production in the lung (Figure 3A-3D). However, Bcl6^fl/fl^CD11c^cre^ mice did not generate lung CD4^+^ T_RM_ cells (Figure 3E-3F) or memory lung T_H_ cells (Figure 3G). Thus, the induction of lung T_FH_ and memory T_H_ cells can be uncoupled. Lung moDCs from Bcl6^fl/fl^CD11c^cre^ mice may selectively lack the ability to induce lung memory T_H_ responses.

**Figure 3.**
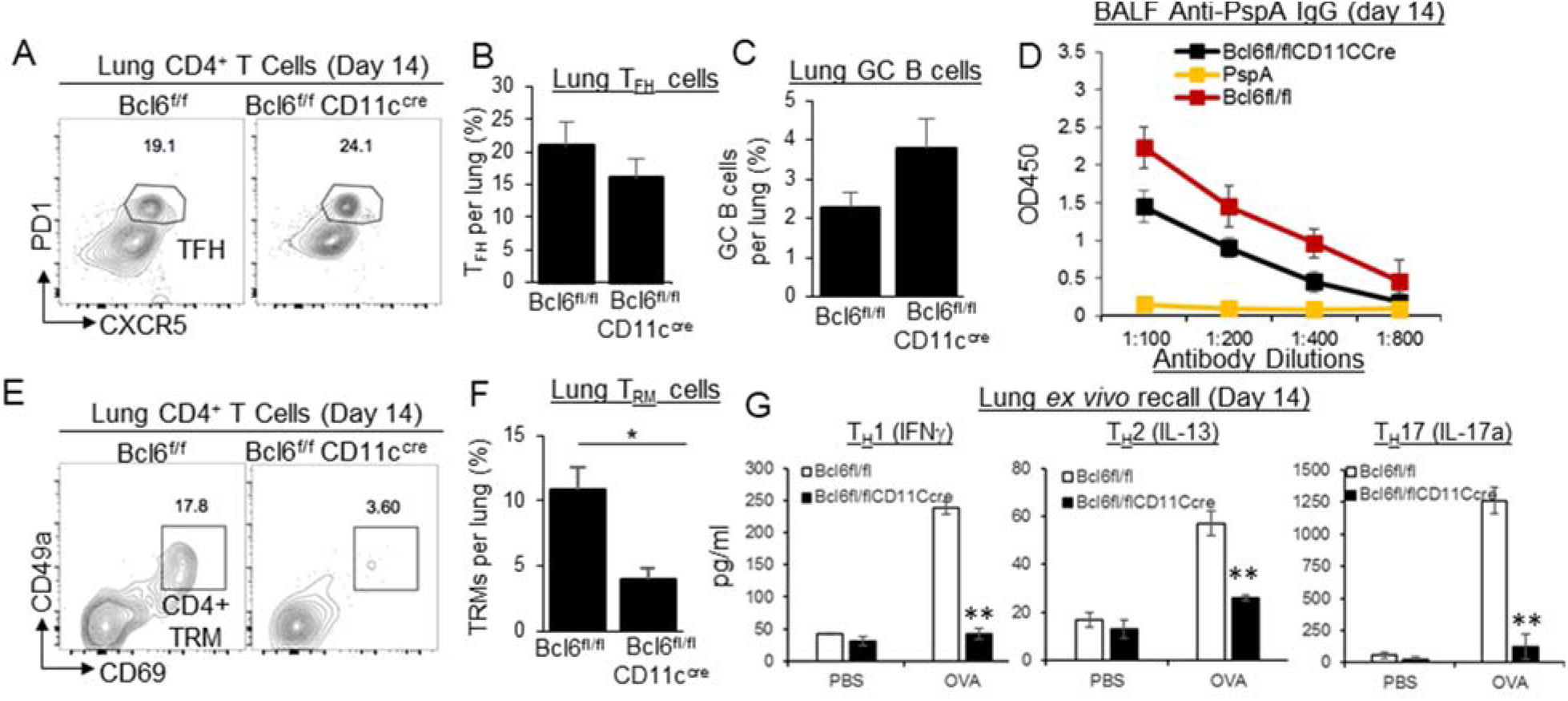
CDG induces lung T_FH_ responses but not CD4^+^ memory T_H_ in Bcl6^fl/fl^CD11c^cre^. **A-C**. Bcl6^fl/fl^ and Bcl6^fl/fl^CD11c^cre^ mice were immunized (*i.n*.) with CDG (5µg) and PspA (2µg). Flow cytometry plots of CD4^+^PD1^+^CXCR5^+^ T_FH_ on day 14 (**A**). Frequency of T_FH_ (**B**) and CD19^+^Bcl6^+^ GC B cells (**C**) in the lung on day 14 post-immunization. (n=3mice/group) Data are representative of three independent experiments. **D**. BALF anti-PspA IgA in mice from (**A)** was determined on day 14 by ELISA. Data are representative of three independent experiments. **E-F**. Flow cytometry plots (**E**) and frequency (**F**) of CD4^+^CD69^+^CD49a^+^ T_RM_ in immunized Bcl6^fl/fl^ and Bcl6^fl/fl^CD11c^cre^ mice from (**A**). Data are representative of three independent experiments. **G**. Lung cells from immunized mice (**A**) were recalled with 5μg/ml PspA for 4 days in culture. Cytokines were measured in the supernatant by ELISA. Data are representative of three independent experiments. Graphs represent the mean with error bars indication s.e.m. *P* values determined by unpaired student *t*-test. \**P*<0.05, \*\**P*<0.001, \*\*\**P*<0.0001.

### HDM induces CD4^+^ memory T_H_ cells in mLNs but not in the lung in Bcl6^fl/fl^CD11c^cre^ mice

Chronic HDM treatment induced lung memory CD4^+^ T cells(Turner et al., 2018) and Bcl6^+^ moDCs (Figure S5B). We found that chronic HDM treatment did not induce lung CD4^+^ T_RM_ cells (Figure 4A-4B) or HDM-specific lung memory T_H_2, T_H_17 responses in the Bcl6^fl/fl^CD11c^cre^ mice (Figure 4C). Chronic HDM treatment still induced strong memory T_H_ responses in mLNs (Figure 4D) and intact serum anti-HDM IgG1 in the Bcl6^fl/fl^CD11c^cre^ mice (Figure 4E), suggesting Bcl6^fl/fl^CD11c^cre^ mice had a selective defective in lung CD4^+^ memory T cell response in chronic HDM treated mice.

**Figure 4.**
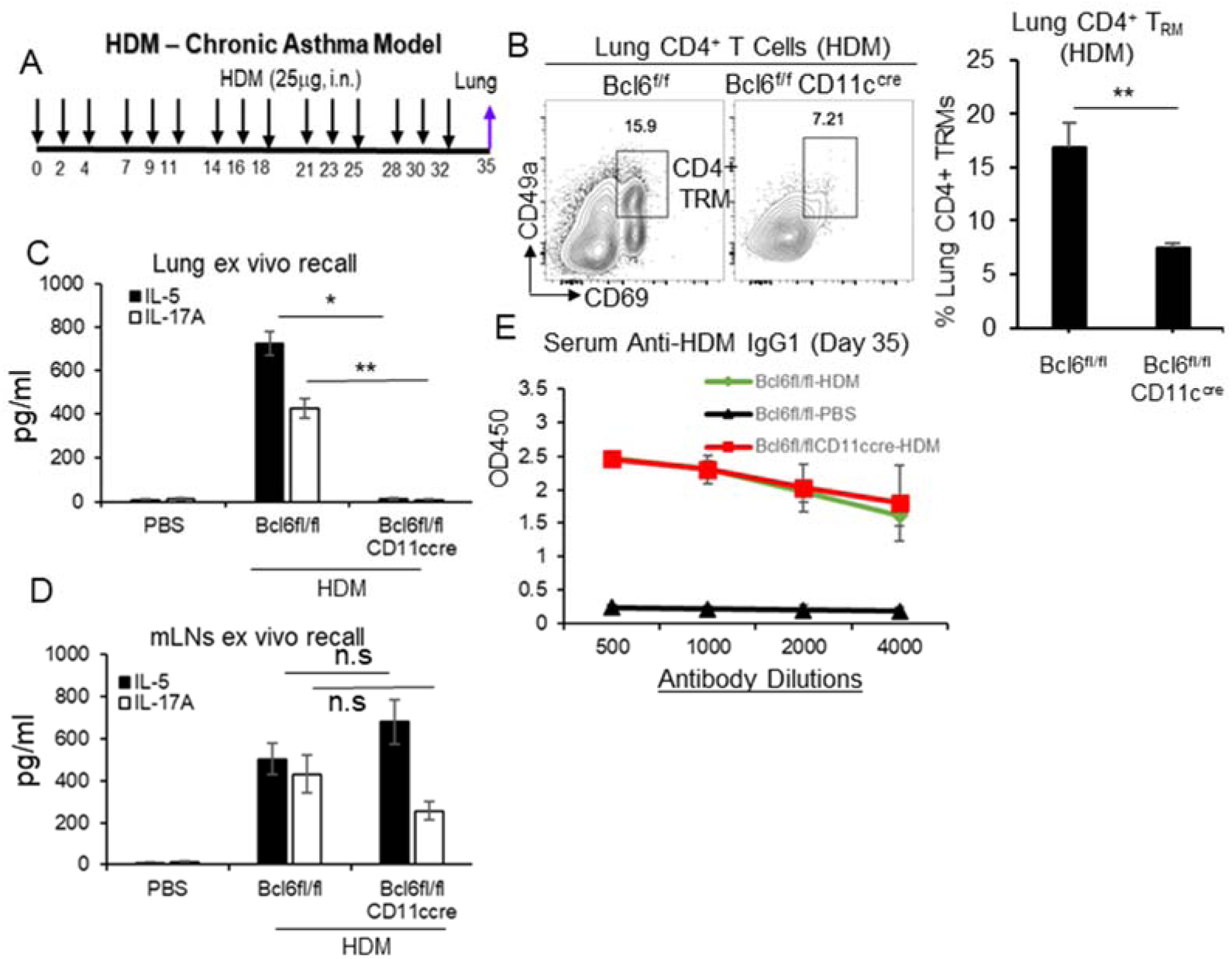
HDM induces CD4^+^ memory T_H_ cells in mLNs but not in the lungs from Bcl6^fl/fl^CD11c^cre^ mice. **A**. A cartoon of chronic HDM treatment in mice. n=3 mice/group. **B**. Flow cytometry analysis (**left**) and frequency (**right**) of CD4^+^CD69^+^CD49a^+^ T_RM_ in HDM-induced Bcl6^fl/fl^ and Bcl6^fl/fl^CD11c^cre^ mice on day 35. (n=3mice/group) Data are representative of three independent experiments. **C-D**. Lung cells (**C**) or mediastinal lymph nodes (mLNs) (**D**) from HDM treated mice on day 35 from (**A**) were stimulated with 25μg/ml HDM for 4 days in culture. T_H_2 and T_H_17 cytokines were measured in the supernatant by ELISA. Data are representative of three independent experiments. **E**. Serum anti-HDM IgG1 were determined in the HDM mice on day 35 from (**A**) by ELISA. Data are representative of three independent experiments. Graphs represent means ± standard error. The significance is determined by one-way ANOVA Tukey’s multiple comparison test (**C, D**) or unpaired Student’s t test (**B**). *p<0.05, \*\**P*<0.001, \*\*\**P*<0.0001.

### Lung moDCs from Bcl6^fl/fl^CD11c^cre^ mice are defective in promoting CD4^+^ memory T_H_ cells in the lung

IL-10 and TGFβ1 are critical cytokines for the induction of T_RM_s(Nath et al., 2019; Thompson et al., 2019). Consistently, we found that on day 14 post-immunization, lung moDCs from immunized Bcl6^fl/fl^CD11C^cre^ mice had decreased production of IL-10 or TGFβ1 (Figure 5A-5B). Lung moDCs from Bcl6^fl/fl^CD11c^cre^ mice also produced less T_H_-promoting cytokines IL-12p70, IL-4, and IL-23 (Figure 5C, S9A-S9B). CCL20 (MIP3A), the CCR6 ligand, recruits lymphocytes and dendritic cells to mucosal lymphoid tissues and is critical for inducing mucosal immune responses(Lee and Korner, 2019). We found that CCL20 production in lung moDCs was also dramatically reduced in CDG immunized Bcl6^fl/fl^CD11c^cre^ mice on day 14 (Figure 5D).

**Figure 5.**
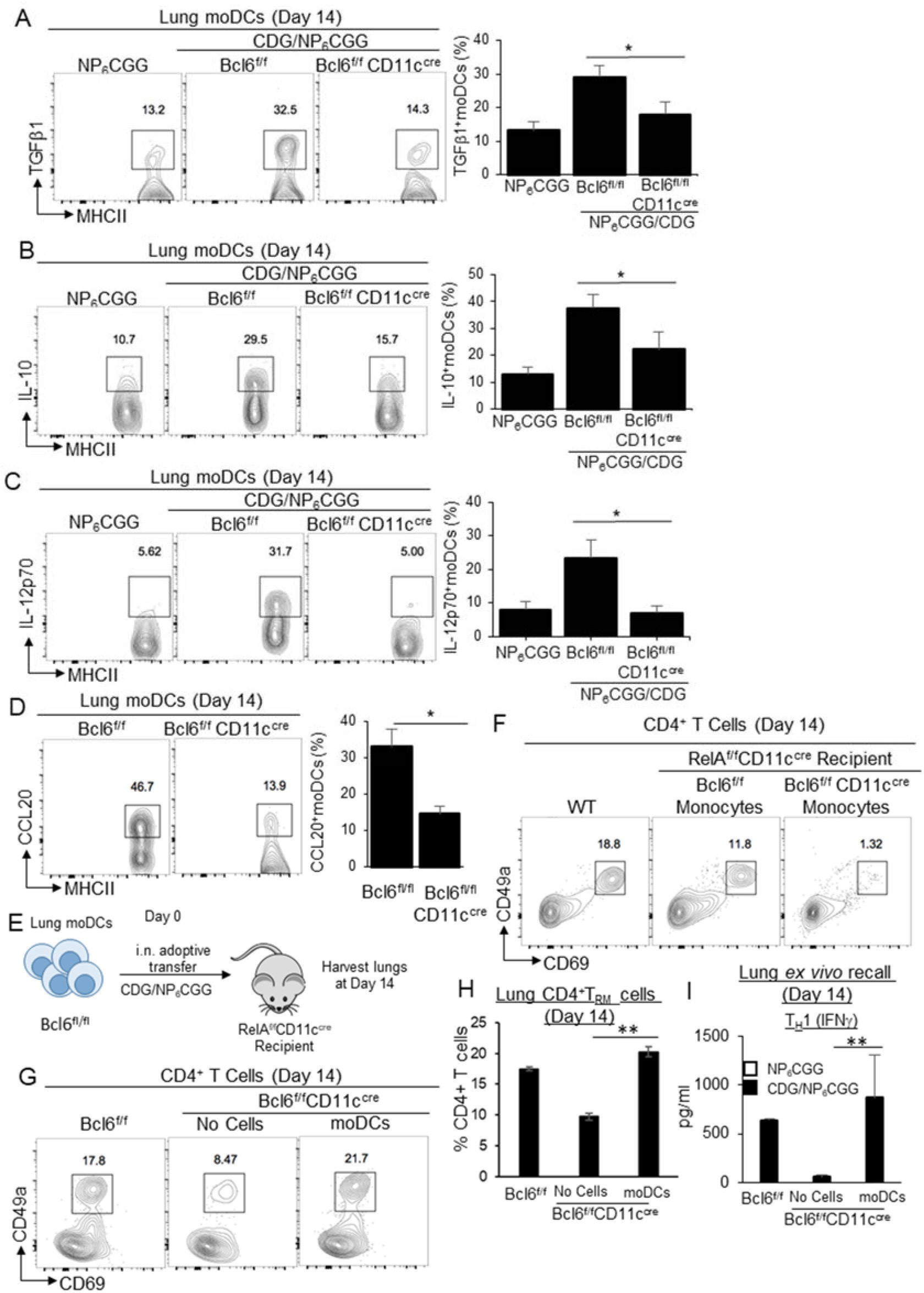
Bcl6^+^ moDCs promote lung mucosal CD4+ memory T_H_ responses. **A-D**. Flow cytometry analysis and frequency of TGFβ1 (**A**), IL-10 (**B**), IL-12p70 (**C**), and CCL20 (**D**) production by lung moDCs in Bcl6^fl/fl^ and Bcl6^fl/fl^CD11c^cre^ mice on day 14 post CDG/NP_6_CGG immunization. (n=3mice/group) Data are representative of two independent experiments. **E**. RelA^fl/fl^CD11c^cre^ mice receiving (*i.n*.) Bcl6^fl/fl^ or Bcl6^fl/fl^CD11c^cre^ bone marrow Ly6C^hi^ monocytes were immunized with CDG/NP_6_CGG. Lung CD4^+^CD69^+^CD49a^+^ T_RM_ cells were determined by Flow cytometry on day 14. (n=3mice/group) Data are representative of two independent experiments. **F**. A cartoon of adoptive moDCs transfer into Bcl6^fl/fl^CD11c^cre^ mice. n=3 mice/group. Sorted naïve WT lung moDCs (∼35,000 cells) were adoptively transferred (*i.n*.) into Bcl6^fl/fl^CD11c^cre^ mice. Recipient Bcl6^fl/fl^CD11c^cre^ mice were immunized with CDG/NP_6_CGG. **G-H**. Lung CD4^+^CD69^+^CD49a^+^ T_RM_ cells were determined by flow cytometry on day 14. **I**. Lung cells from immunized mice were recalled with 5μg/ml NP_6_CGG for 4 days in culture. Cytokines were measured in the supernatant by ELISA. (n=3mice/group) Data are representative of two independent experiments. Graphs represent means ± standard error. The significance is determined by one-way ANOVA Tukey’s multiple comparison test (**A, B, C, H, I**) or unpaired Student’s t test (**D**). * p<0.05, \*\**P*<0.001, \*\*\**P*<0.0001.

To further demonstrate the lung moDCs from Bcl6^fl/fl^CD11c^cre^ mice can not induce lung memory T_H_ responses, we did the adoptive cell transfer experiments in RelA^fl/fl^CD11c^cre^ and Bcl6^fl/fl^CD11c^cre^ mice. First, we isolated Ly6C^hi^ monocytes from Bcl6^fl/fl^ and Bcl6^fl/fl^CD11c^cre^ mice and transferred (*i.n*.) them into CDG/OVA immunized RelA^fl/fl^CD11c^cre^ mice on day 0. The moDCs in RelA^fl/fl^CD11c^cre^ can not be activated due to the lack of RelA (Figure 1). On day 14, we examined the lung CD4^+^ T_RM_ cells. Only the RelA^fl/fl^CD11c^cre^ mice receiving monocytes from Bcl6^fl/fl^, not Bcl6^fl/fl^CD11c^cre^ mice generated lung CD4^+^ T_RM_ cells (Figure 5E), suggesting monocytes or monocytes-derived cells from Bcl6^fl/fl^CD11c^cre^ mice were defective in inducing CD4^+^ T_RM_ cells in the lung.

Monocytes can differentiate into macrophages as well. To exclude the possibility of monocyte-derived macrophages in lung memory CD4^+^ T cells response, we adoptively transferred moDCs. We sorted out lung moDCs from a naïve C57BL/6 mouse and transferred lung moDCs into (*i.n*.) immunized Bcl6^fl/fl^CD11c^cre^ mice (Figure 5F). We examined lung memory T_H_ responses on day 14 post-immunization. Again, WT moDCs restored memory T_H_1 responses and lung CD4^+^ T_RM_ cells in the immunized Bcl6^fl/fl^CD11c^cre^ (Figure 5G-5I). Together, the data indicated that lung moDCs from Bcl6^fl/fl^CD11c^cre^ mice can not promote CD4^+^ memory T_H_ responses in the lung and it is moDCs cell-intrinsic defect.

### Lung TNFR2^+^ cDC2 secrete TNF to generate Bcl6^+^ moDCs in the lung

The data so far concluded the functional heterogeneity of lung moDCs. The data also suggested that Bcl6 could be a marker to separate moDCs subpopulations that promote memory T_H_ vs. T_FH_ cells respectively. The Bcl6^+^ moDCs produce T_H_ and T_RM_ promoting cytokines, chemokines and are likely responsible for lung memory T_H_ responses. We previously showed that the lung TNFR2^+^ cDC2 population induces lung memory T_H_ responses(Mansouri et al., 2019). Furthermore, moDCs did not take up intranasal administered CDG(Mansouri et al., 2019). We, thus, hypothesized that the CDG activated TNFR2^+^ cDC2, which drives moDCs differentiation into Bcl6^+^ moDCs.

IRF4^fl/fl^CD11c^cre^ mice lack the cDC2 population and do not respond to CDG immunization(Mansouri et al., 2019). We adoptively transferred WT lung TNFR2^+^ cDC2 into IRF4^fl/fl^CD11c^cre^ mice and immunized the recipient mice with CDG/OVA. We examined lung Bcl6^+^ moDCs in the recipient mice on day 14 post-immunization. As expected, IRF4^fl/fl^CD11c^cre^ received PBS did not generate Bcl6^+^ moDCs. In contrast, IRF4^fl/fl^CD11c^cre^ received WT TNFR2^+^ cDC2 population had lung Bcl6^+^ moDCs (Figure 6A).

**Figure 6.**
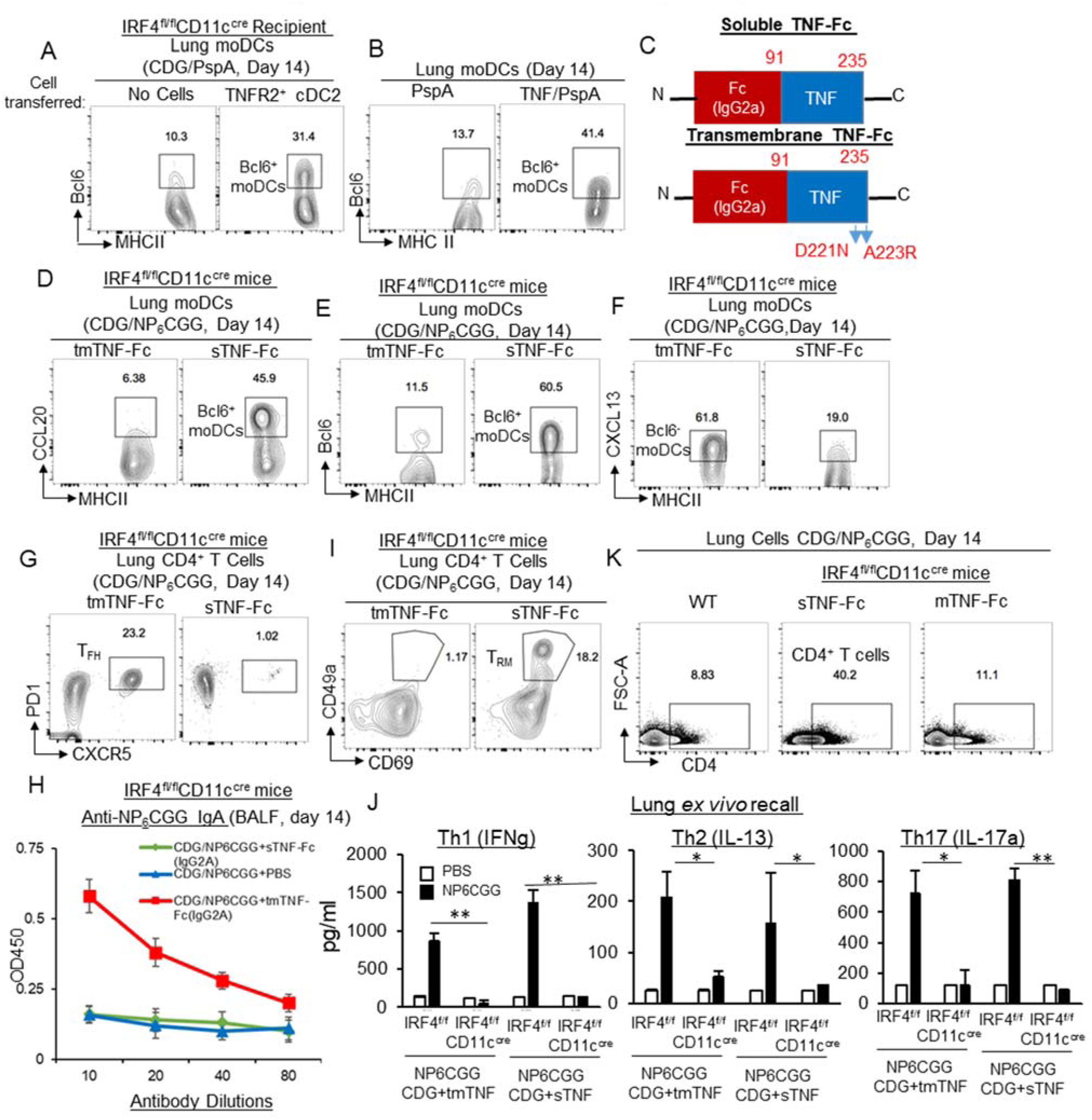
Targeting soluble and transmembrane TNF to moDCs *in vivo* generate functionally distinct lung Bcl6^+^ and Bcl6^-^ moDCs respectively. **A**. Sorted naïve lung TNFR2^+^ cDC2 were adoptively transferred (*i.n*.) into IRF4^fl/fl^CD11c^cre^ mice and immunized with CDG/PspA. On Day 14, Bcl6^+^ moDCs were determined by flow cytometry. (n=3 mice/group) Data are representative of three independent experiments. **B**. WT mice were treated (*i.n*.) with 200ng recombinant TNF and PspA or PspA alone. On day 14, Bcl6^+^ moDCs in the lung were determined by Flow cytometry. (n=3 mice/group) Data are representative of two independent experiments. **C**. Cartoon illustrating the sTNF-Fc (IgG2A) and tmTNF-Fc (IgG2A) fusion proteins. **D-F**. IRF4^fl/fl^CD11c^cre^ mice were immunized (*i.n*.) with CDG/NP_6_CGG and 100ng tmTNF-Fc (IgG2A) or 100ng sTNF-Fc (IgG2A). Lungs were harvested on day 14. Flow cytometry analysis of CCL20 (**D**), Bcl6 (**E**), and CXCL13 (**F**) expression by lung moDCs. (n=3 mice/group) Data are representative of three independent experiments. **G-H**. Flow cytometry analysis of lung T_FH_ (**G**) and lung antigen-specific IgA (**H**) in mice from (**D-F**). Lungs were harvested on Day 14. Data are representative of three independent experiments. **I**. Flow cytometry analysis of lung CD4^+^ T_RM_ cells in mice from **(D-F)** on day 14. Data are representative of three independent experiments. **J**. Lung cells from mice in **(D-F)** on day 14 post-immunization were recalled with 5μg/ml NP_6_CGG in culture for 4 days. Cytokines were measured in the supernatant by ELISA. Data are representative of three independent experiments. **K**. Flow cytometry analysis of lung CD4^+^ T cells in mice from **(D-F)** on day 14. Data are representative of three independent experiments. Graphs represent means ± standard error. The significance is represented by one-way ANOVA Tukey’s multiple comparison test. * p<0.05, \*\**P*<0.001, \*\*\**P*<0.0001.

How did the lung TNFR2^+^ cDC2 population stimulate the differentiation of Bcl6^+^ moDCs in the lung? We previously discovered that lung TNFR2^-^ cDC2 population stimulated moDCs maturation by the transmembrane TNF - TNFR2 interaction(Mansouri et al., 2019). TNF is essential for CDG mucosal adjuvant activity(Blaauboer et al., 2014). cDC2 is the main source for intranasal CDG-induced lung TNF(Mansouri et al., 2019). We hypothesize that TNFR^+^ cDC2 cells promote the differentiation of Bcl6^+^ moDCs via TNF secretion. Indeed, the intranasal administration of recombinant soluble TNF/PspA induced lung Bcl6^+^ moDCs in C57BL/6J mice on day 14 post-immunization (Figure 6B).

### CDG/TNF-Fc(IgG2A) and CDG/TNF_D221N/A223R_-Fc(IgG2A) fusion proteins promote the generation of lung Bcl6^+^ and Bcl6^-^ moDCs respectively in the IRF4^fl/fl^CD11c^cre^ mice

To demonstrate that lung TNF acts on moDCs directly to promote the differentiation of Bcl6^+^ moDCs, we generated a fusion protein TNF-Fc (IgG2A) to target TNF to moDCs (Figure 6C). moDCs express the high-affinity FcR, FcγRI, also known as CD64 that is not found on cDCs or lymphocytes(Langlet et al., 2012). FcγRI bind the Fc of IgG2a with the highest affinity (10^8^M^-1^), more than 1,000 fold higher than its next binding partner IgG2b-Fc(Guilliams et al., 2014). Indeed, intranasal administration of APC-conjugated mouse IgG2A was taken up exclusively by CD64^+^ lung cells, including MHC II^hi^ CD64^+^ moDCs (Figure S10). The CD64^+^ MHC class II^low/int^ macrophages were also targeted. However, macrophages are dispensable for CDG mucosal adjuvanticity *in vivo*(Blaauboer et al., 2015; Mansouri et al., 2019).

We used IRF4^fl/fl^CD11c^cre^ because CDG does not generate lung TNF in the IRF4^fl/fl^CD11c^cre(Mansouri et al., 2019)^, which facilitates the TNF-Fc (IgG 2A) complement experiment. CDG immunization generates soluble TNF and transmembrane TNF (tmTNF)(Mansouri et al., 2019). We fused TNF with the Fc portion of IgG2A to generate TNF-Fc (IgG2A). We also made a TNF_D221N/A223R_-Fc (IgG2A) fusion protein that targets transmembrane TNF to moDCs. The TNF_D221N/A223R_ mutant mimics transmembrane TNF that binds only to TNFR2, not TNFR1(Loetscher et al., 1993).

We immunized IRF4^fl/fl^CD11c^cre^ mice with CDG/NP_6_CGG with tmTNF-Fc (IgG2A) or TNF-Fc (IgG2A). On day 14, we examined lung moDCs. The addition of TNF-Fc (IgG2A), not TNF_D221N/A223R_-Fc (IgG2A), generated Bcl6^+^ and CCL20^+^ moDCs in the lung (Figure 6D-6E). In contrast, the addition of TNF_D221N/A223R_-Fc (IgG2A), not TNF-Fc (IgG2A) treatment, generated CXCL13^+^ moDCs (Figure 6F). Importantly, TNF_D221N/A223R_-Fc (IgG2A), not TNF-Fc (IgG2A) administration, generated lung T_FH_ and lung IgA in IRF4^fl/fl^CD11c^cre^ mice (Figure 6G-6H).

### TNF-Fc (IgG2A) generated CD4^+^ T_RM_-like cells in the lung of the IRF4^fl/fl^CD11c^cre^ mice

IRF4^fl/fl^CD11c^cre^ mice lack cDC2 including the migratory TNFR2^+^ cDCc2 that generates T_EF_ cells in the dLNs(Mansouri et al., 2019). We hypothesized that lung moDCs provide “Signal 4” to convert T_EF_ cells into memory CD4^+^ T cells in the lung (Figure 7). Thus, we were surprised to find that CDG/TNF-Fc (IgG2A) administration generated CD4^+^ T_RM_–like cells in the lung in IRF4^fl/fl^CD11c^cre^ mice (Figure 6I). We suspected that these CD4^+^ T_RM_–like (CD49a^+^CD69^+^) cells in IRF4^fl/fl^CD11c^cre^ mice may not be antigen-specific lung memory cells. We did the *ex vivo* recall assay. Indeed, neither TNF_D221N/A223R_-Fc (IgG2A) nor TNF-Fc (IgG2A) restored lung memory T_H_ responses in CDG immunized IRF4^fl/fl^CD11c^cre^ mice (Figure 6J). Thus, TNF-Fc (IgG2A) did not generate NP_6_CGG-specific lung CD4^+^ memory T cells in CDG/NP_6_CGG immunized IRF4^fl/fl^CD11c^cre^ mice, albeit these T_RM_-like cells could be lung memory CD4^+^ T cells against unknown antigens.

**Figure 7.**
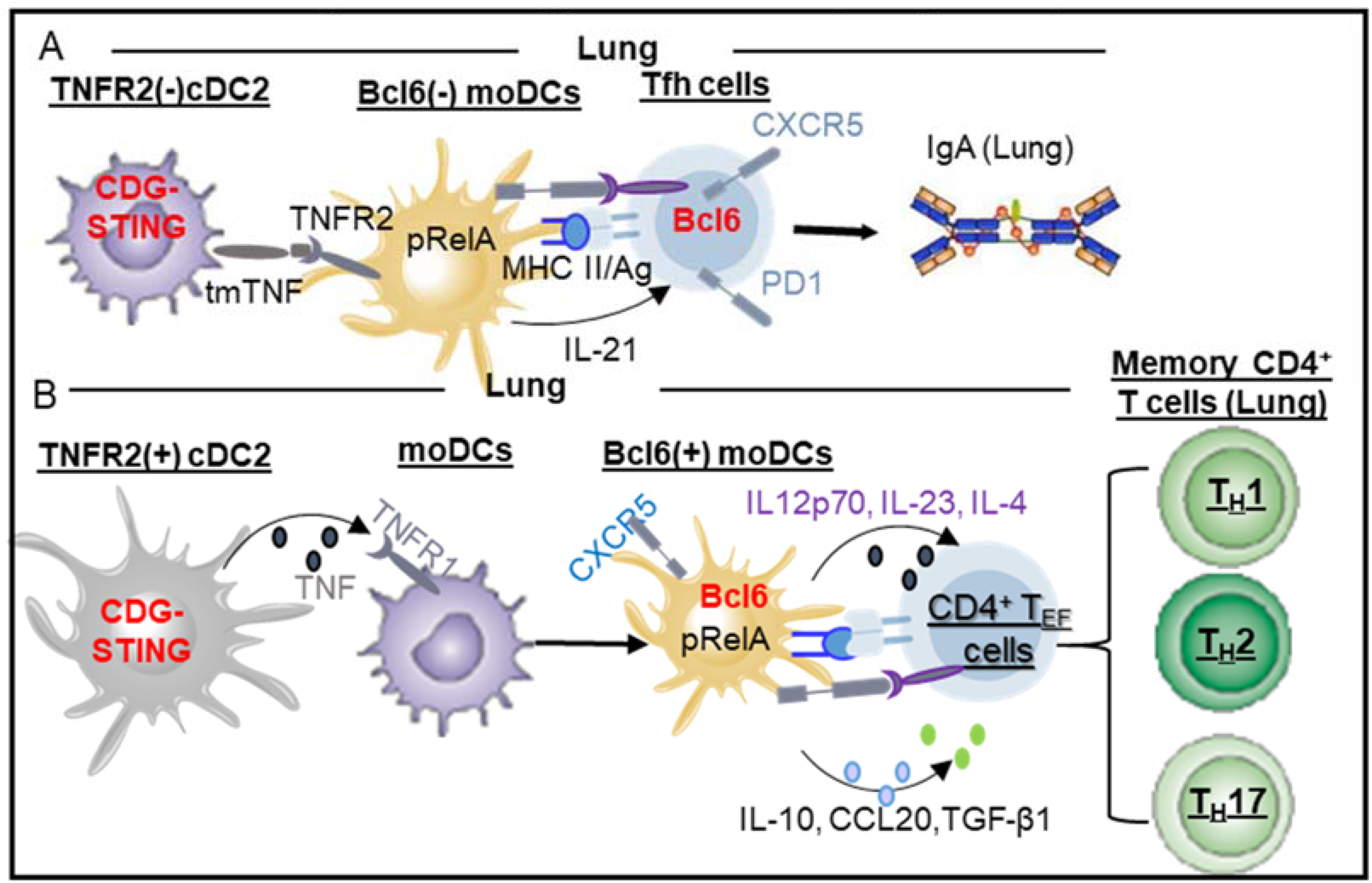
Model – Differentiated lung Bcl6^+^ and Bcl6^-^ moDCs induce CD4^+^ memory T_H_ cells and T_FH_ cells in the lung respectively. **A**. We previously showed(Mansouri et al., 2019) that intranasal administration of adjuvant CDG activates STING pathway in TNFR2(-) cDC2 population to mature moDCs via the transmembrane TNF and generate T_FH_ cells, IgA in lung mucosa. **B**. CDG activates STING pathway in the lung TNFR2(+) cDC2 population to secrete TNF(Mansouri et al., 2019), which differentiate moDCs into Bcl6^+^ moDCs that provide “Signal 4” in lung mucosa to drive CD4^+^ T_EF_ cells into lung memory CD4^+^ T_H_ cells.

We noticed that TNF-Fc (IgG2A), not TNF_D221N/A223R_-Fc (IgG2A), dramatically increased numbers of lung CD4^+^ T cells in CDG immunized IRF4^fl/fl^CD11c^cre^ mice (Figure 6K), likely due to the CCL20 production by TNF-Fc (IgG2A) (Figure 6D). Lung Bcl6^+^ moDCs might prime these infiltrating lung CD4^+^ T cells to generate these CD4^+^ T_RM_-like cells in the lung of IRF4^fl/fl^CD11c^cre^ mice. Altogether, the moDCs targeting TNF fusion protein data strongly argued that moDCs can differentiate into functionally distinct Bcl6^+^ and Bcl6^-^ moDCs in the lung via the stimulation by soluble or transmembrane TNF.

## Discussion

moDCs have a universal presence at the site of inflammation and can promote CD4^+^ and CD8^+^ T cells responses *in vivo*. Yet, we know very little about how moDCs achieve this plasticity *in vivo*. Consequently, we have few approaches to control this common and versatile APC population *in vivo* during inflammation. The most exciting discoveries in this report were i) the identification of functionally distinct moDCs subpopulations responsible for lung CD4^+^ memory T_H_ and T_FH_ cells respectively; ii) the development of moDCs-targeting TNF fusion proteins to control lung moDCs differentiation *in vivo*.

We used the mucosal adjuvant CDG system to study moDCs *in vivo*. CDG activates mainly the STING pathway(Blaauboer et al., 2014) and does not induce lung damage(Blaauboer et al., 2015). Its simplicity uncouples the lung mucosal vs. systemic, lung memory T_H_ vs. lung T_FH_ vaccine responses, thus facilitates the extraction of the underlying mechanisms. The discovery from the CDG study was confirmed in the inflammatory HDM-treated mice. We showed that similar to the CDG-induced vaccine adjuvant responses, Bcl6^fl/fl^CD11c^cre^ mice lack HDM-induced lung memory CD4^+^ T_H_ cells while retained mLNs memory CD4^+^ T_H_ responses. Lung TNF generates Bcl6^+^ moDCs. TNF is a hallmark of inflammation. We propose that Bcl6^+^ moDCs exist at the site of all inflammation and may play a critical role in driving T_H_ cells-mediated inflammatory responses.

We separated differentiated lung moDCs based on their expression of Bcl6. We proposed that lung moDCs differentiate into Bcl6^+^ moDCs to promote lung memory T_H_ responses. First, the Bcl6^+^ moDCs are likely a heterogeneous population as well promoting T_H_1, T_H_2 or T_H_17 responses in the lung. Second, though Bcl6 expression in lung moDCs is likely functional relevant, we did not establish a causal link between Bcl6 expression in Bcl6^+^ moDCs and their ability to promote memory CD4^+^ T_H_ responses. Both conditional Bcl6 mice had DCs developmental defects. The Bcl6^fl/fl^LysM^cre^ mice do not have functional moDCs while the Bcl6^fl/fl^CD11c^cre^ mice had no cDC1. Nevertheless, CDG did not induce Bcl6 expression in lung moDCs from the Bcl6^fl/fl^CD11c^cre^ mice on day 14 post-immunization. LysM and CD11c-driven Bcl6 conditional mice have been described before(Zhu et al., 2019). However, the defect in lung moDCs were not reported(Zhu et al., 2019). Here, we established a critical role of Bcl6 in lung moDCs development. Future study is needed to understand the cell-intrinsic role of Bcl6 in controlling the function of the differentiated Bcl6^+^ lung moDCs.

How Bcl6^+^ moDCs promote the induction of lung CD4^+^ memory T cells? Lung CD4^+^ T cell memory formation depends on re-engagement of CD4^+^ T cells with APC at their effector stage (Signal 4) (Bautista et al., 2016; McKinstry et al., 2014). The lung Bcl6^+^ moDCs are a differentiated moDCs population appeared after the initial priming. We propose that the differentiated Bcl6^+^ moDCs are the long-thought lung APCs that provide the “Signal 4” during the effector stage to induce lung memory CD4^+^ T cells. Indeed, lung Bcl6^+^ moDCs bore antigens and expressed CD80, CD86 on day 14 post-immunization. However, mature Bcl6^-^ moDCs exist during the same timeframe and present antigen in the lung on day 14 post-immunization. In fact, the Bcl6^-^ moDCs induced lung T_FH_ cells. So why does Bcl6^-^ mature moDCs can not drive lung CD4^+^ memory T_H_ cell induction? In the lung, most CD4^+^ T_RM_ cells are in B cell follicles(Hondowicz et al., 2016; Shinoda et al., 2012; Takamura et al., 2016; Turner et al., 2014). Bcl6 expression upregulates CXCR5 on moDCs. CXCR5^+^ DCs were found to localize near B cell follicles in the marginal zone of the spleen and the dermis of the skin(Saeki et al., 2000; Yu et al., 2002). We hypothesize that the expression of chemokine receptor CXCR5 positions the mature Bcl6^+^ moDCs on the boundary of the B-T cell zone, where they provide “Signal 4” to incoming CD4^+^ T_EF_ cells via the high endothelial venules and promote CD4^+^ memory cells formation.

Recall response is a hallmark of immune memory cells, which establishes antigen-specificity and distinguish T_H_1, T_H_2, and T_H_17 memory CD4^+^ T cells. We used lung *ex vivo* recall assay to determine the total lung CD4^+^ memory T cells, including memory T_H_1, T_H_2, and T_H_17 cells induced by immunization. We reasoned that though the lung memory T_H_ responses are mainly from lung CD4^+^ T_RM_ cells, T_EM_ cells or other unknown memory CD4 T cells in the lung may also contribute to the total lung memory T_H_ responses(Reagin and Klonowski, 2018). Furthermore, growing evidence suggest that T_RM_ cells, even at the same tissue site, *e.g*. lung, are heterogeneous(Mueller and Mackay, 2016). In fact, we identified CD4^+^ T_RM_-like cells in TNF-Fc (IgG2A) adjuvanted CDG immunized IRF4^fl/fl^CD11c^cre^ mice, but these T_RM_-like cells could not be recalled by the immunized antigen. Our goal is to generate antigen-specific protection against lung infections for which all memory CD4^+^ T cells in the lung contribute. We believe measuring total lung memory T_H_1/2/17 cells by *ex vivo* lung recall is a better indicator for the vaccine efficacy in lung mucosa than enumerating total cell numbers of lung CD69^+^CD49a^+^ CD4 T_RM_ cells.

In summary, moDCs can differentiate into Bcl6^+^ and Bcl6^-^ lung moDCs to promote CD4^+^ T_H_ cells and T_FH_ cells, respectively in the lung mucosa. Targeted activation of moDCs by soluble and tmTNF fusion proteins could be a valuable method to enhance lung T-cells memory responses.

## Methods

### KEY RESOURCES TABLE

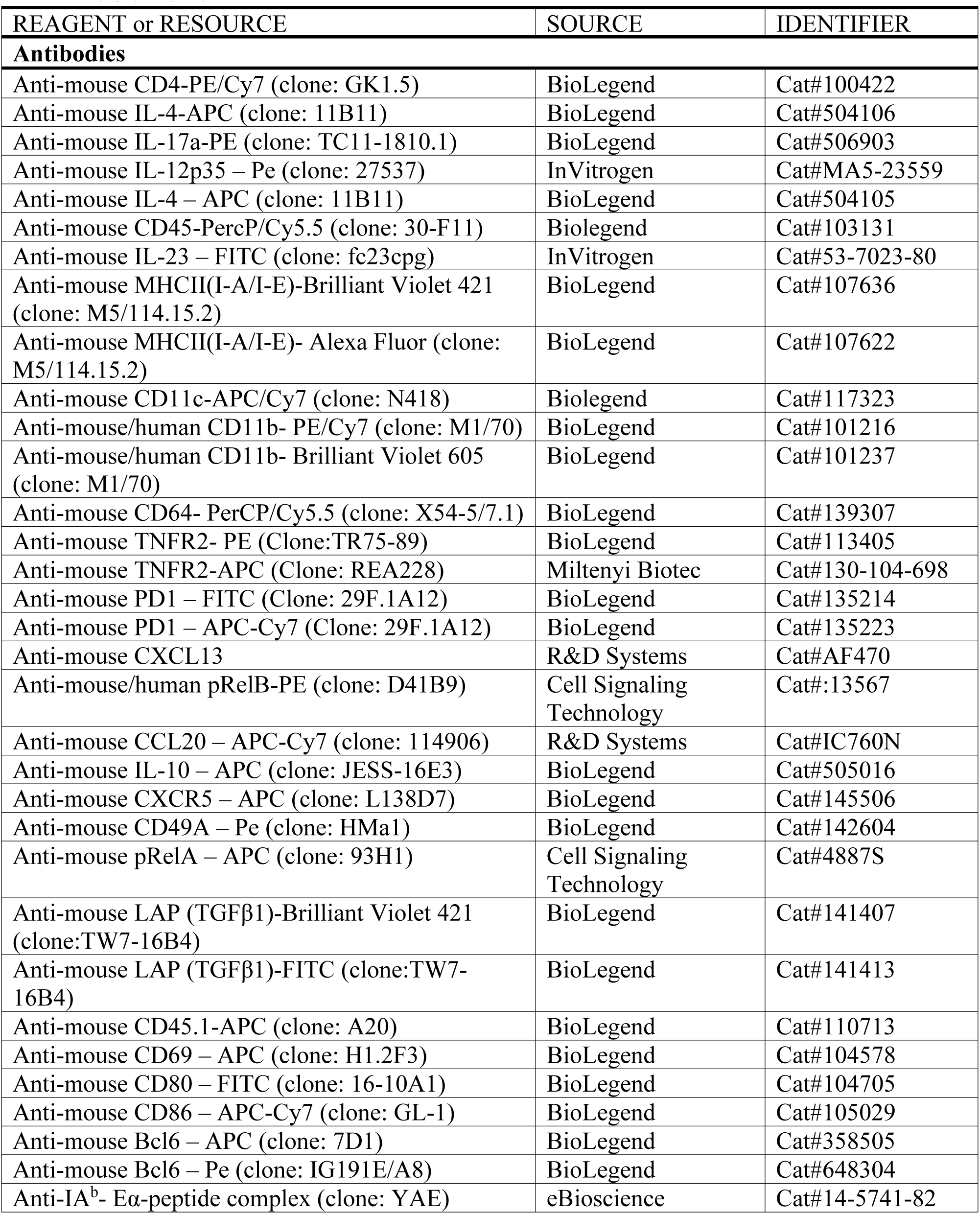
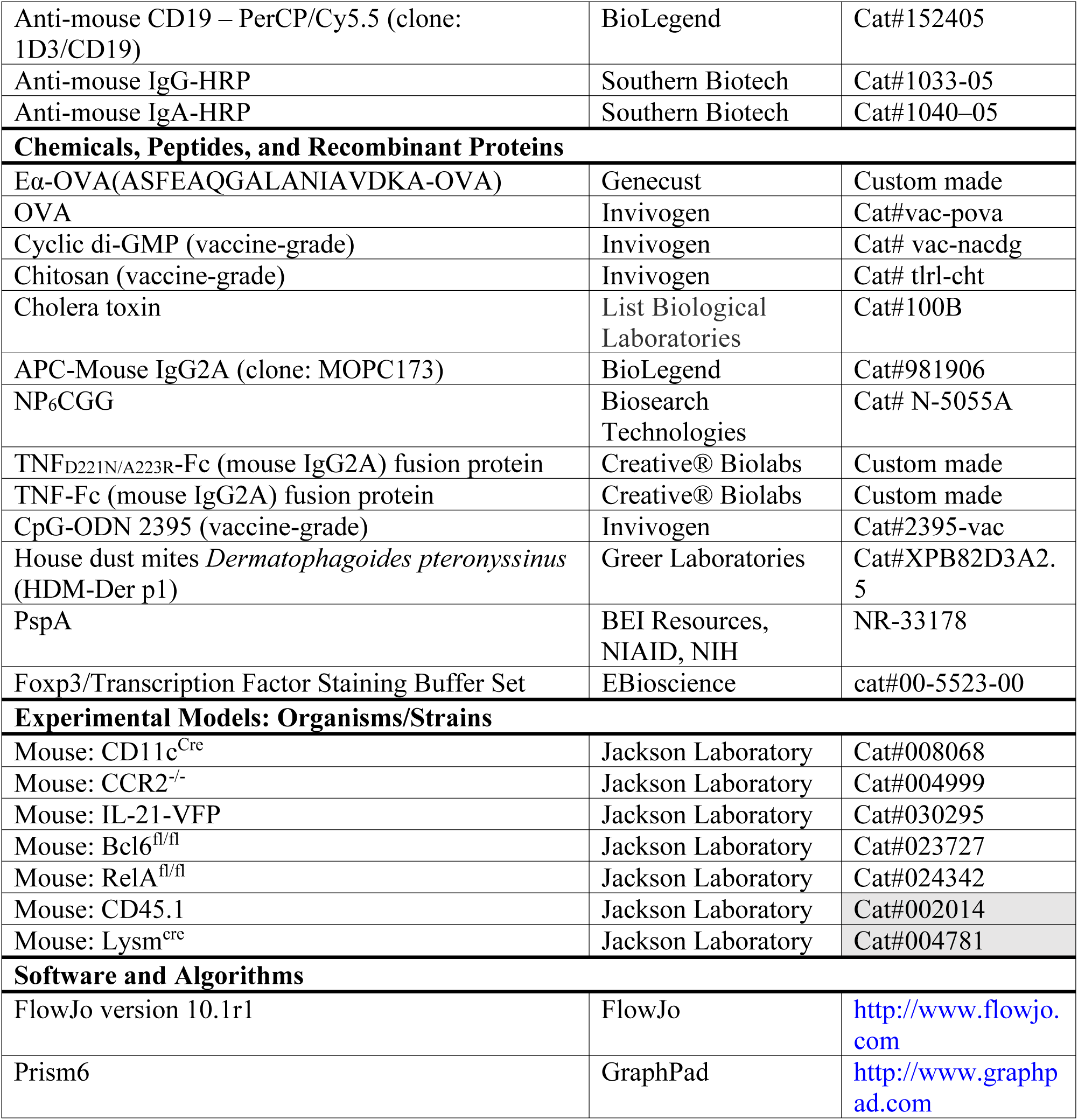

### Mice

Age- and gender-matched mice (2 - 3 months old) were used for the initial immunization or HDM treatment. C57BL/6J, B6.CD45.1(#002014), *Bcl6*^*fl/fl*^ (#023727), *CCR2*^*-/-*^ (#004999), *RelA*^*fl/fl*^ (#024342), *IL-21-VFP(#030295), CD11c*^*cre*^ (#008068) and *LysM*^*cre*^ (#004781) mice were purchased from The Jackson Laboratory. Mice were housed and bred under pathogen-free conditions in the Animal Research Facility at the University of Florida. All mouse experiments were performed by the regulations and approval of the Institutional Animal Care and Use Committee at the University of Florida, IACUC number 201909362.

### Reagents

Eα-OVA (ASFEAQGALANIAVDKA-OVA) fusion protein was produced by GeneCust. TNF-IgG2a-Fc fusion proteins were produced by Creative® Biolabs. WT mouse TNF (aa80∼235) or mutant mouse TNF (TNF_D221N/A223R_) was fused with Fc portion of the mouse IgG2A. Following reagents were from Invivogen, endotoxin-free OVA (vac-pova), vaccine-grade CDG (vac-nacdg), vaccine-grade CpG-ODN 2395 (2395-vac-1), chitosan (tlrl-cht). House dust mite (XPB82D3A2.5, D. pter) was from Stallergenes Greer. Cholera toxin was from List Biological Laboratories (100B). Following reagents were from eBioscience, YAE mAb (14-5741-82). Following reagents are from Biosearch Technologies, NP_6_CGG (N-5055A).

The following reagent was obtained through BEI Resources, NIAID, NIH: Streptococcus pneumoniae Family 2, Clade 3 Pneumococcal Surface Protein A (PspA UAB099) with C-Terminal Histidine Tag, Recombinant from Escherichia coli, NR-33179.

### Intranasal CDG Immunization

Groups of mice were intranasally vaccinated with CDG (5µg) adjuvanted antigen (2µg) or antigen alone (Blaauboer et al., 2014). For intranasal vaccination, animals were anesthetized using isoflurane in an E-Z Anesthesia system (Euthanex Corp, Palmer, PA). Antigen, with or without CDG was administered in 30µl saline. Sera were collected at the indicated time points after the last immunization. The antigen-specific Abs were determined by ELISA. Secondary Abs used were anti-mouse IgG-HRP (Southern Biotech, 1033– 05), and anti-mouse IgA-HRP (Southern Biotech, 1040–05).To determine Ag-specific T_H_ response, splenocytes and lung cells from antigen or CDG + antigen immunized mice were stimulated with 5µg/ml antigen for four days in culture. T_H_ cytokines were measured in the supernatant by ELISA.

### HDM-induced chronic asthma

House dust mites *Dermatophagoides pteronyssinus* (HDM-Der p1, Greer Laboratories, cat no. XPB82D3A2.5) was suspended in endotoxin-free PBS at a concentration of 5mg/ml. To induce chronic asthma, mice were challenged with 25µg HDM every other day for five consecutive weeks. To measure HDM-specific lung memory T_H_, lung cells were restimulated with 25µg/ml of HDM for 4 days. Memory T_H_ cells were measured in the supernatant by ELISA.

### Isolation of lung cells

Cells were isolated from the lung, as previously described(Mansouri et al., 2019). The lungs were perfused with ice-cold PBS and removed. Lungs were digested in DMEM containing 200μg/ml DNase I (Roche, 10104159001), 25μg/ml Liberase TM (Roche, 05401119001) at 37°C for 2 hours. Red blood cells were then lysed and a single cell suspension was prepared by filtering through a 70-µm cell strainer.

### Intracellular staining

For transcription factor Bcl6 staining of murine and human cells, cells were fixed and permeabilized with the Foxp3 staining buffer set (eBioscience, cat no 00-5523-00). The intracellular cytokine staining was performed using the Cytofix/Cytoperm™ kit from BD Biosciences (cat#555028). The single lung cell suspension was fixed in Cytofix/perm buffer (BD Biosciences) in the dark for 20min at RT. Fixed cells were then washed and kept in Perm/Wash buffer at 4°C. Golgi-plug was present during every step before fixation.

### Mouse monocyte purification and adoptive transfer

Mouse Ly6C^hi^ monocytes were purified from the bone marrow of naïve mice (C57BL/6J or CD45.1) following the protocol according to the manufacturer (Stemcell Technologies, 19861). Mice were intranasally vaccinated with CDG (5µg) and antigen. Ly6C^hi^ monocytes (1.5 million/mouse) were administered intranasally into immunized mice at 30mins, 2hrs and 4hrs post-immunization.

### Flow cytometry

Single-cell suspensions were stained with fluorescent-dye-conjugated antibodies in PBS containing 2% FBS and 1mM EDTA. Surface stains were performed at 4°C for 20 min. Cells were washed and stained with surface markers. Cells were then fixed and permeabilized (eBioscience, cat no. 00-5523-00) for intracellular cytokine stain. Data were acquired on a BD LSRFortessa and analyzed using the FlowJo software package (FlowJo, LLC). Cell sorting was performed on the BD FACSAriaIII Flow Cytometer and Cell Sorter.

### Experimental Design

Data exclusion was justified when positive or negative control did not work. All experiments will be repeated at least three times. All repeats are biological replications that involve the same experimental procedures on different mice. Experiments comparing different genotypes, adjuvant responses are designed with individual treatments being assigned randomly. Where possible, treatments will be assigned blindly to the experimenter by another individual in the lab. When comparing samples from different groups, samples from each group will be analyzed in concert, thereby preventing any biases that may arise from analyzing individual treatments on different days

### Statistical Analysis

All data are expressed as means ± SEM. Statistical significance was evaluated using Prism 6.0 software. One-way ANOVA was performed with post hoc Tukey’s multiple comparison test, Mann-Whitney U-test, or Student’s t-test applied as appropriate for comparisons between groups. A p-value of <0.05 was considered significant.

## Supplemental Material

A pdf file containing 10 Supplementary Figures can be found online.

## Acknowledgments

This work was supported by NIH grants AI110606, AI125999, AI132865 (to L.J.).

## Author contributions

S.M. and L.J. conceived and designed the research. S.M., D.S.K, H.G performed experiments and analyzed the data. L.J. helped with experiments and analysis. S.M and L.J. wrote the manuscript. L.J. supervised the research.

## References

Allen, A.C., Wilk, M.M., Misiak, A., Borkner, L., Murphy, D., and Mills, K.H.G. (2018). Sustained protective immunity against Bordetella pertussis nasal colonization by intranasal immunization with a vaccine-adjuvant combination that induces IL-17-secreting TRM cells. Mucosal Immunol 11, 1763–1776.

Bautista, B.L., Devarajan, P., McKinstry, K.K., Strutt, T.M., Vong, A.M., Jones, M.C., Kuang, Y., Mott, D., and Swain, S.L. (2016). Short-Lived Antigen Recognition but Not Viral Infection at a Defined Checkpoint Programs Effector CD4 T Cells To Become Protective Memory. J Immunol 197, 3936–3949.

Blaauboer, S.M., Gabrielle, V.D., and Jin, L. (2014). MPYS/STING-mediated TNF-alpha, not type I IFN, is essential for the mucosal adjuvant activity of (3’-5’)-cyclic-di-guanosine-monophosphate in vivo. J Immunol 192, 492–502.

Blaauboer, S.M., Mansouri, S., Tucker, H.R., Wang, H.L., Gabrielle, V.D., and Jin, L. (2015). The mucosal adjuvant cyclic di-GMP enhances antigen uptake and selectively activates pinocytosis-efficient cells in vivo. Elife 4.

Casey, K.A., Fraser, K.A., Schenkel, J.M., Moran, A., Abt, M.C., Beura, L.K., Lucas, P.J., Artis, D., Wherry, E.J., Hogquist, K., et al. (2012). Antigen-independent differentiation and maintenance of effector-like resident memory T cells in tissues. J Immunol 188, 4866–4875.

Chow, K.V., Sutherland, R.M., Zhan, Y., and Lew, A.M. (2017). Heterogeneity, functional specialization and differentiation of monocyte-derived dendritic cells. Immunol Cell Biol 95, 244–251.

Desai, P., Tahiliani, V., Stanfield, J., Abboud, G., and Salek-Ardakani, S. (2018). Inflammatory monocytes contribute to the persistence of CXCR3(hi) CX3CR1(lo) circulating and lung-resident memory CD8(+) T cells following respiratory virus infection. Immunol Cell Biol 96, 370–378.

Dunbar, P.R., Cartwright, E.K., Wein, A.N., Tsukamoto, T., Tiger Li, Z.R., Kumar, N., Uddback, I.E., Hayward, S.L., Ueha, S., Takamura, S., and Kohlmeier, J.E. (2019). Pulmonary monocytes interact with effector T cells in the lung tissue to drive TRM differentiation following viral infection. Mucosal Immunol.

Ebensen, T., Schulze, K., Riese, P., Link, C., Morr, M., and Guzman, C.A. (2007). The bacterial second messenger cyclic diGMP exhibits potent adjuvant properties. Vaccine 25, 1464–1469.

Guilliams, M., Bruhns, P., Saeys, Y., Hammad, H., and Lambrecht, B.N. (2014). The function of Fcgamma receptors in dendritic cells and macrophages. Nat Rev Immunol 14, 94–108.

Haddadi, S., Vaseghi-Shanjani, M., Yao, Y., Afkhami, S., D’Agostino, M.R., Zganiacz, A., Jeyanathan, M., and Xing, Z. (2019). Mucosal-Pull Induction of Lung-Resident Memory CD8 T Cells in Parenteral TB Vaccine-Primed Hosts Requires Cognate Antigens and CD4 T Cells. Front Immunol 10, 2075.

Hondowicz, B.D., An, D., Schenkel, J.M., Kim, K.S., Steach, H.R., Krishnamurty, A.T., Keitany, G.J., Garza, E.N., Fraser, K.A., Moon, J.J., et al. (2016). Interleukin-2-Dependent Allergen-Specific Tissue-Resident Memory Cells Drive Asthma. Immunity 44, 155–166.

Iborra, S., Martinez-Lopez, M., Khouili, S.C., Enamorado, M., Cueto, F.J., Conde-Garrosa, R., Del Fresno, C., and Sancho, D. (2016). Optimal Generation of Tissue-Resident but Not Circulating Memory T Cells during Viral Infection Requires Crosspriming by DNGR-1(+) Dendritic Cells. Immunity 45, 847–860.

Ko, H.J., Brady, J.L., Ryg-Cornejo, V., Hansen, D.S., Vremec, D., Shortman, K., Zhan, Y., and Lew, A.M. (2014). GM-CSF-responsive monocyte-derived dendritic cells are pivotal in Th17 pathogenesis. J Immunol 192, 2202–2209.

Krawczyk, C.M., Shen, H., and Pearce, E.J. (2007). Memory CD4 T cells enhance primary CD8 T-cell responses. Infect Immun 75, 3556–3560.

Kumar, B.V., Ma, W., Miron, M., Granot, T., Guyer, R.S., Carpenter, D.J., Senda, T., Sun, X., Ho, S.H., Lerner, H., et al. (2017). Human Tissue-Resident Memory T Cells Are Defined by Core Transcriptional and Functional Signatures in Lymphoid and Mucosal Sites. Cell Rep 20, 2921–2934.

Langlet, C., Tamoutounour, S., Henri, S., Luche, H., Ardouin, L., Gregoire, C., Malissen, B., and Guilliams, M. (2012). CD64 expression distinguishes monocyte-derived and conventional dendritic cells and reveals their distinct role during intramuscular immunization. J Immunol 188, 1751–1760.

Lee, A.Y.S., and Korner, H. (2019). The CCR6-CCL20 axis in humoral immunity and T-B cell immunobiology. Immunobiology 224, 449–454.

Lees, J.R., and Farber, D.L. (2010). Generation, persistence and plasticity of CD4 T-cell memories. Immunology 130, 463–470.

Leon, B., Lopez-Bravo, M., and Ardavin, C. (2007). Monocyte-derived dendritic cells formed at the infection site control the induction of protective T helper 1 responses against Leishmania. Immunity 26, 519–531.

Loetscher, H., Stueber, D., Banner, D., Mackay, F., and Lesslauer, W. (1993). Human tumor necrosis factor alpha (TNF alpha) mutants with exclusive specificity for the 55-kDa or 75-kDa TNF receptors. J Biol Chem 268, 26350–26357.

Mackay, L.K., Stock, A.T., Ma, J.Z., Jones, C.M., Kent, S.J., Mueller, S.N., Heath, W.R., Carbone, F.R., and Gebhardt, T. (2012). Long-lived epithelial immunity by tissue-resident memory T (TRM) cells in the absence of persisting local antigen presentation. Proc Natl Acad Sci U S A 109, 7037–7042.

Mansouri, S., Patel, S., Katikaneni, D.S., Blaauboer, S.M., Wang, W., Schattgen, S., Fitzgerald, K., and Jin, L. (2019). Immature lung TNFR2(-) conventional DC 2 subpopulation activates moDCs to promote cyclic di-GMP mucosal adjuvant responses in vivo. Mucosal Immunol 12, 277–289.

McKinstry, K.K., Strutt, T.M., Bautista, B., Zhang, W., Kuang, Y., Cooper, A.M., and Swain S.L. (2014). Effector CD4 T-cell transition to memory requires late cognate interactions that induce autocrine IL-2. Nat Commun 5, 5377.

McKinstry, K.K., Strutt, T.M., Kuang, Y., Brown, D.M., Sell, S., Dutton, R.W., and Swain, S.L. (2012). Memory CD4+ T cells protect against influenza through multiple synergizing mechanisms. J Clin Invest 122, 2847–2856.

McKinstry, K.K., Strutt, T.M., and Swain, S.L. (2010). The potential of CD4 T-cell memory. Immunology 130, 1–9.

McMaster, S.R., Wein, A.N., Dunbar, P.R., Hayward, S.L., Cartwright, E.K., Denning, T.L., and Kohlmeier, J.E. (2018). Pulmonary antigen encounter regulates the establishment of tissue-resident CD8 memory T cells in the lung airways and parenchyma. Mucosal Immunol 11, 1071–1078.

Mueller, S.N., and Mackay, L.K. (2016). Tissue-resident memory T cells: local specialists in immune defence. Nat Rev Immunol 16, 79–89.

Nath, A.P., Braun, A., Ritchie, S.C., Carbone, F.R., Mackay, L.K., Gebhardt, T., and Inouye, M. (2019). Comparative analysis reveals a role for TGF-beta in shaping the residency-related transcriptional signature in tissue-resident memory CD8+ T cells. PLoS One 14, e0210495.

Plantinga, M., Guilliams, M., Vanheerswynghels, M., Deswarte, K., Branco-Madeira, F., Toussaint, W., Vanhoutte, L., Neyt, K., Killeen, N., Malissen, B., et al. (2013). Conventional and monocyte-derived CD11b(+) dendritic cells initiate and maintain T helper 2 cell-mediated immunity to house dust mite allergen. Immunity 38, 322–335.

Randolph, G.J., Beaulieu, S., Lebecque, S., Steinman, R.M., and Muller, W.A. (1998). Differentiation of monocytes into dendritic cells in a model of transendothelial trafficking. Science 282, 480–483.

Reagin, K.L., and Klonowski, K.D. (2018). Incomplete Memories: The Natural Suppression of Tissue-Resident Memory CD8 T Cells in the Lung. Front Immunol 9, 17.

Saeki, H., Wu, M.T., Olasz, E., and Hwang, S.T. (2000). A migratory population of skin-derived dendritic cells expresses CXCR5, responds to B lymphocyte chemoattractant in vitro, and co-localizes to B cell zones in lymph nodes in vivo. Eur J Immunol 30, 2808–2814.

Sathaliyawala, T., Kubota, M., Yudanin, N., Turner, D., Camp, P., Thome, J.J., Bickham, K.L., Lerner, H., Goldstein, M., Sykes, M., et al. (2013). Distribution and compartmentalization of human circulating and tissue-resident memory T cell subsets. Immunity 38, 187–197.

Schreiner, D., and King, C.G. (2018). CD4+ Memory T Cells at Home in the Tissue: Mechanisms for Health and Disease. Front Immunol 9, 2394.

Serbina, N.V., Salazar-Mather, T.P., Biron, C.A., Kuziel, W.A., and Pamer, E.G. (2003). TNF/iNOS-producing dendritic cells mediate innate immune defense against bacterial infection. Immunity 19, 59–70.

Sharma, M.D., Rodriguez, P.C., Koehn, B.H., Baban, B., Cui, Y., Guo, G., Shimoda, M., Pacholczyk, R., Shi, H., Lee, E.J., et al. (2018). Activation of p53 in Immature Myeloid Precursor Cells Controls Differentiation into Ly6c(+)CD103(+) Monocytic Antigen-Presenting Cells in Tumors. Immunity 48, 91–106 e106.

Shinoda, K., Tokoyoda, K., Hanazawa, A., Hayashizaki, K., Zehentmeier, S., Hosokawa, H., Iwamura, C., Koseki, H., Tumes, D.J., Radbruch, A., and Nakayama, T. (2012). Type II membrane protein CD69 regulates the formation of resting T-helper memory. Proc Natl Acad Sci U S A 109, 7409–7414.

Smith, N.M., Wasserman, G.A., Coleman, F.T., Hilliard, K.L., Yamamoto, K., Lipsitz, E., Malley, R., Dooms, H., Jones, M.R., Quinton, L.J., and Mizgerd, J.P. (2018). Regionally compartmentalized resident memory T cells mediate naturally acquired protection against pneumococcal pneumonia. Mucosal Immunol 11, 220–235.

Stockinger, B., Bourgeois, C., and Kassiotis, G. (2006). CD4+ memory T cells: functional differentiation and homeostasis. Immunol Rev 211, 39–48.

Strutt, T.M., McKinstry, K.K., Dibble, J.P., Winchell, C., Kuang, Y., Curtis, J.D., Huston, G., Dutton, R.W., and Swain, S.L. (2010). Memory CD4+ T cells induce innate responses independently of pathogen. Nat Med 16, 558-564, 551p following 564.

Takamura, S., Yagi, H., Hakata, Y., Motozono, C., McMaster, S.R., Masumoto, T., Fujisawa, M., Chikaishi, T., Komeda, J., Itoh, J., et al. (2016). Specific niches for lung-resident memory CD8+ T cells at the site of tissue regeneration enable CD69-independent maintenance. J Exp Med 213, 3057–3073.

Teijaro, J.R., Turner, D., Pham, Q., Wherry, E.J., Lefrancois, L., and Farber, D.L. (2011). Cutting edge: Tissue-retentive lung memory CD4 T cells mediate optimal protection to respiratory virus infection. J Immunol 187, 5510–5514.

Thompson, E.A., Darrah, P.A., Foulds, K.E., Hoffer, E., Caffrey-Carr, A., Norenstedt, S., Perbeck, L., Seder, R.A., Kedl, R.M., and Lore, K. (2019). Monocytes Acquire the Ability to Prime Tissue-Resident T Cells via IL-10-Mediated TGF-beta Release. Cell Rep 28, 1127–1135 e1124.

Turner, D.L., Bickham, K.L., Thome, J.J., Kim, C.Y., D’Ovidio, F., Wherry, E.J., and Farber, D.L. (2014). Lung niches for the generation and maintenance of tissue-resident memory T cells. Mucosal Immunol 7, 501–510.

Turner, D.L., Goldklang, M., Cvetkovski, F., Paik, D., Trischler, J., Barahona, J., Cao, M., Dave, R., Tanna, N., D’Armiento, J.M., and Farber, D.L. (2018). Biased Generation and In Situ Activation of Lung Tissue-Resident Memory CD4 T Cells in the Pathogenesis of Allergic Asthma. J Immunol 200, 1561–1569.

Wakim, L.M., Smith, J., Caminschi, I., Lahoud, M.H., and Villadangos, J.A. (2015). Antibody-targeted vaccination to lung dendritic cells generates tissue-resident memory CD8 T cells that are highly protective against influenza virus infection. Mucosal Immunol 8, 1060–1071.

Wilk, M.M., and Mills, K.H.G. (2018). CD4 TRM Cells Following Infection and Immunization: Implications for More Effective Vaccine Design. Front Immunol 9, 1860.

Yang, L., Yu, Y., Kalwani, M., Tseng, T.W., and Baltimore, D. (2011). Homeostatic cytokines orchestrate the segregation of CD4 and CD8 memory T-cell reservoirs in mice. Blood 118, 3039–3050.

Yu, P., Wang, Y., Chin, R.K., Martinez-Pomares, L., Gordon, S., Kosco-Vibois, M.H., Cyster, J., and Fu, Y.X. (2002). B cells control the migration of a subset of dendritic cells into B cell follicles via CXC chemokine ligand 13 in a lymphotoxin-dependent fashion. J Immunol 168, 5117–5123.

Zhang, T.T., Liu, D., Calabro, S., Eisenbarth, S.C., Cattoretti, G., and Haberman, A.M. (2014). Dynamic expression of BCL6 in murine conventional dendritic cells during in vivo development and activation. PLoS One 9, e101208.

Zhu, B., Zhang, R., Li, C., Jiang, L., Xiang, M., Ye, Z., Kita, H., Melnick, A.M., Dent, A.L., and Sun, J. (2019). BCL6 modulates tissue neutrophil survival and exacerbates pulmonary inflammation following influenza virus infection. Proc Natl Acad Sci U S A 116, 11888–11893.

