## Supplemental Figures for "Mucosal vaccine adjuvant cyclic di-GMP differentiates lung moDCs into Bcl6^+^ and Bcl6^−^ mature moDCs to induce lung memory CD4^+^ T_H_ cells and lung T_FH_ cells respectively"

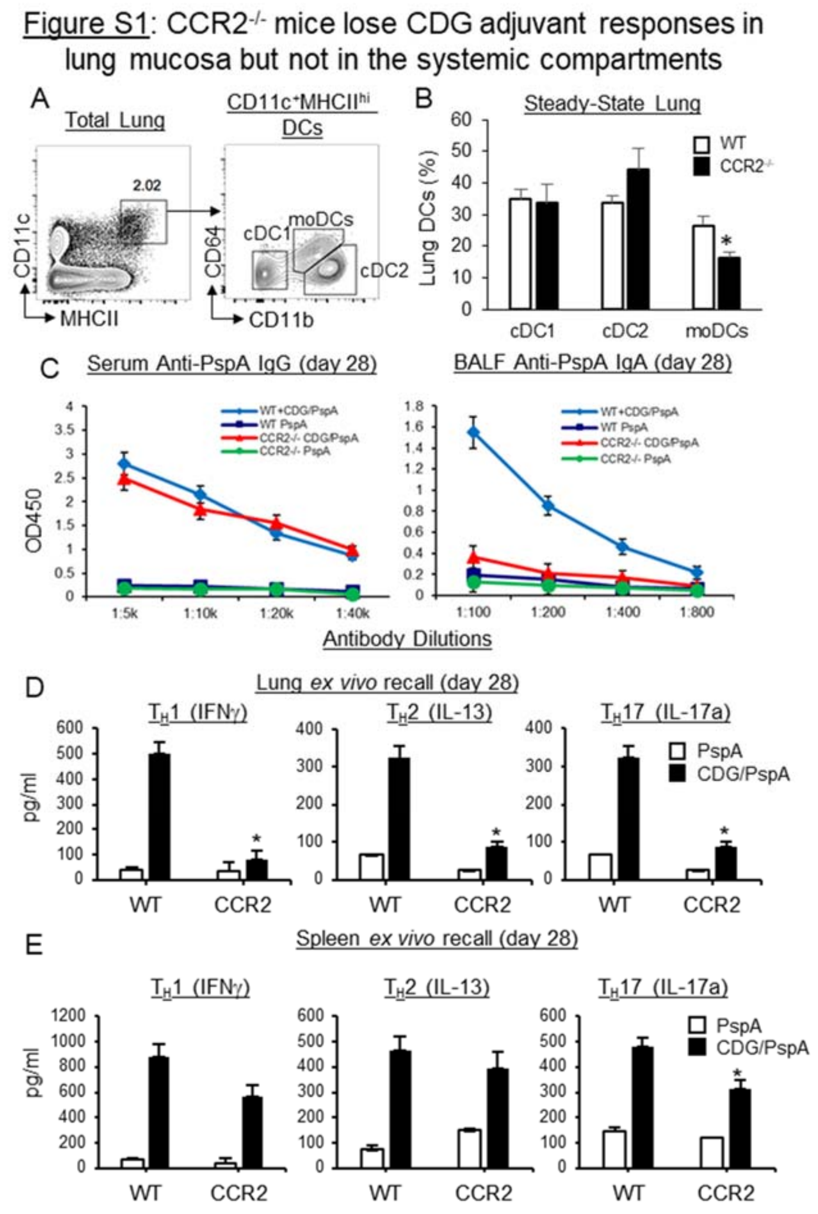


**Figure S1. CCR2$^{-/-}$ mice selectively lose CDG adjuvant responses in lung mucosa but not in the systemic compartments. A.** Gating strategy for lung DCs. cDC1 are MHCII$^{hi}$CD11c$^{+}$CD11b$^{-}$CD64$^{-}$, moDCs are MHCII$^{hi}$CD11c$^{+}$CD11b$^{+}$CD64$^{+}$, and cDC2 are MHCII$^{hi}$CD11c$^{+}$CD11b$^{+}$CD64$^{-}$ **B.** Frequency of lung DCs at steady-state in WT and CCR2$^{-/-}$ mice. (n=3mice/group). Data are representative of four independent experiments. **C.** WT and CCR2$^{-/-}$ mice were immunized (*i.n.*) with two doses (14 days apart) of PspA or PspA plus CDG (5μg). Anti-PspA IgG in serum (left)

and IgA in BALF (right) were determined by ELISA 28 days post-immunization. (n=3mice/group). Data are representative of three independent experiments. **D-E.** Lung cells (**D**) or splenocytes (**E**) from immunized mice (**C**) were recalled with 5μg/ml PspA for 4 days in culture. Cytokines were measured in the supernatant by ELISA. Data are representative of three independent experiments. Graphs represent the mean with error bars indication s.e.m. *P* values determined by unpaired student *t*-test (**B**) or one-way ANOVA Tukey's multiple comparison test (**D, E**). $*P<0.05$

Figure S2: CDG adjuvant-induced lung mucosal $T_{FH}$, GC B cells and $CD4^+$ TRM cells were impaired in $RelA^{fl/fl}CD11c^{cre}$ mice

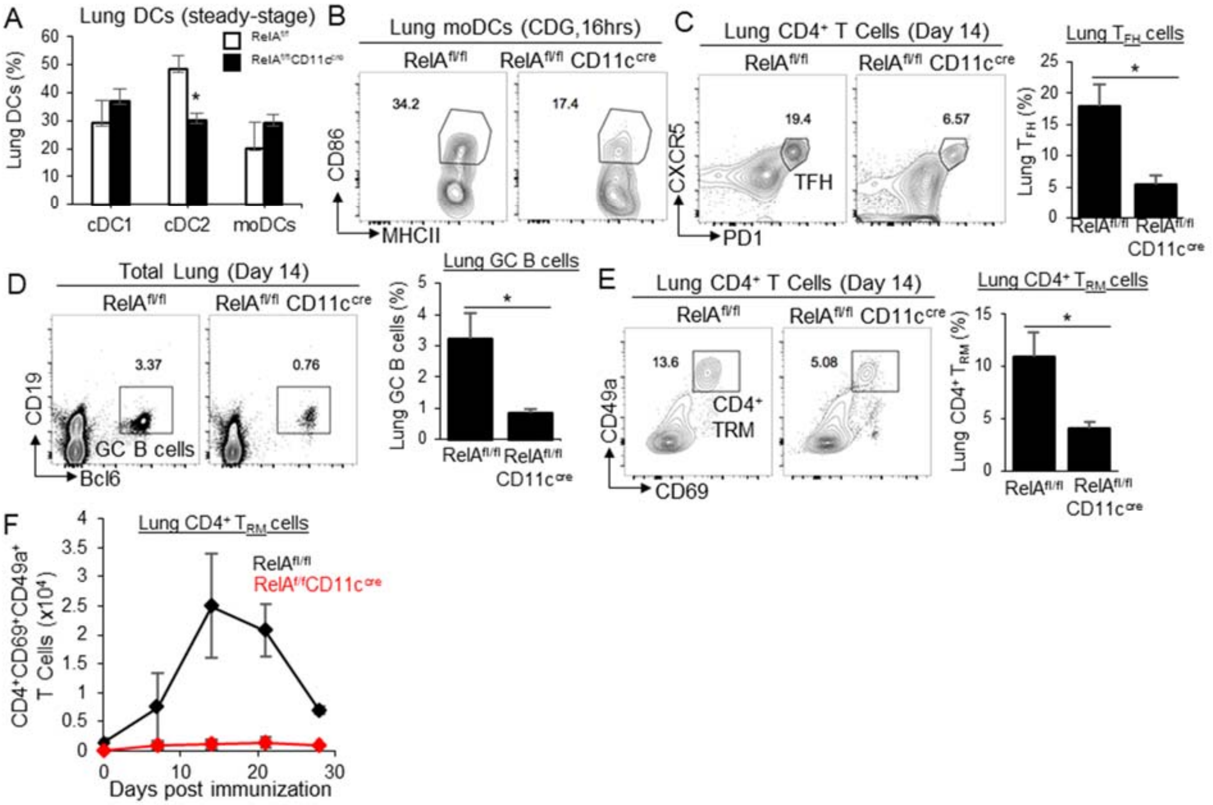


**Figure S2. CDG adjuvant-induced lung mucosal $T_{FH}$, GC B cells and $CD4^+$ TRM cells were impaired in $RelA^{fl/fl}CD11c^{cre}$ mice. A.** Frequency of lung DCs at steady-state in $RelA^{fl/fl}$ and $RelA^{fl/fl}CD11c^{cre}$ mice. (n=3mice/group) Data are representative of two independent experiments. **B.** Flow cytometry analysis of $CD86^+$ moDCs in $RelA^{fl/fl}$ and $RelA^{fl/fl}CD11c^{cre}$ mice administered (*i.n.*) with 5μg CDG for 16 hours. (n=3mice/group). Data are representative of three independent experiments. **C-E.** $RelA^{fl/fl}$ and $RelA^{fl/fl}CD11c^{cre}$ mice were immunized (*i.n.*) with CDG (5μg) and PspA (2μg). Flow cytometry plots of lung $CD4^+PD1^+CXCR5^+$ $T_{FH}$ (**C**), $CD4^+CD69^+CD49a^+$ $T_{RM}$ (**E**) and lung $CD19^+Bcl6^+$ B cells (**D**) on day 14. (n=3mice/group). Data are representative of three independent experiments. **F.** The kinetics of lung $CD4^+CD69^+CD49a^+$ $T_{RM}$ in $RelA^{fl/fl}$ and $RelA^{fl/fl}CD11c^{cre}$ mice at indicated time points after immunization (*i.n.*) with CDG (5μg) and PspA

(2μg). (n=3mice/group) Data are representative of three independent experiments. Graphs represent the mean with error bars indication s.e.m. *P* values determined by unpaired student *t*-test. $*P<0.05$

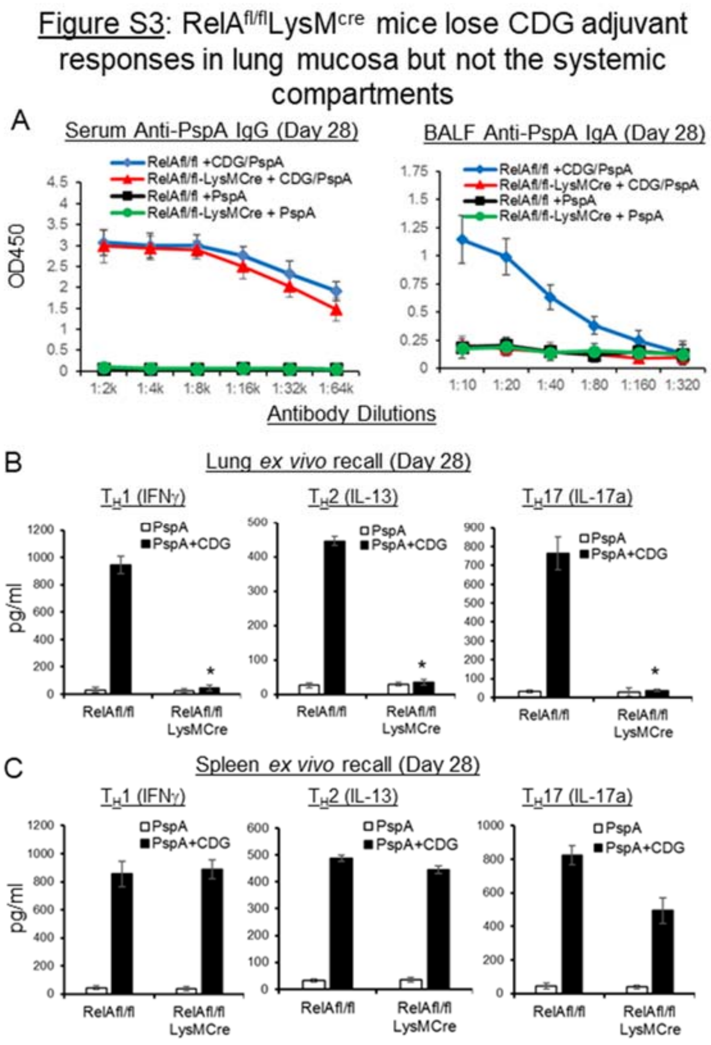


**Figure S3. RelA$^{fl/fl}$LysM$^{cre}$ mice selectively lose CDG adjuvant responses in the lung mucosa but not in the systemic compartments. A.** RelA$^{fl/fl}$ and RelA$^{fl/fl}$LysM$^{cre}$ mice were immunized (*i.n.*) with two doses (14 days apart) of PspA or PspA plus CDG (5μg). Anti-PspA IgG in serum (left) and IgA in BALF (right) were determined by ELISA 28 days post-immunization. (n=3mice/group) Data are representative of two independent experiments. **B-C.** Lung cells (**B**) or splenocytes (**C**) from immunized RelA$^{fl/fl}$ and RelA$^{fl/fl}$LysM$^{cre}$ mice (**A**) were recalled with 5μg/ml PspA for 4 days in culture. Cytokines were measured in the supernatant by ELISA. Graphs represent the mean with error bars indication s.e.m. *P* values determined by one-way ANOVA Tukey's multiple comparison test. *$P$<0.05

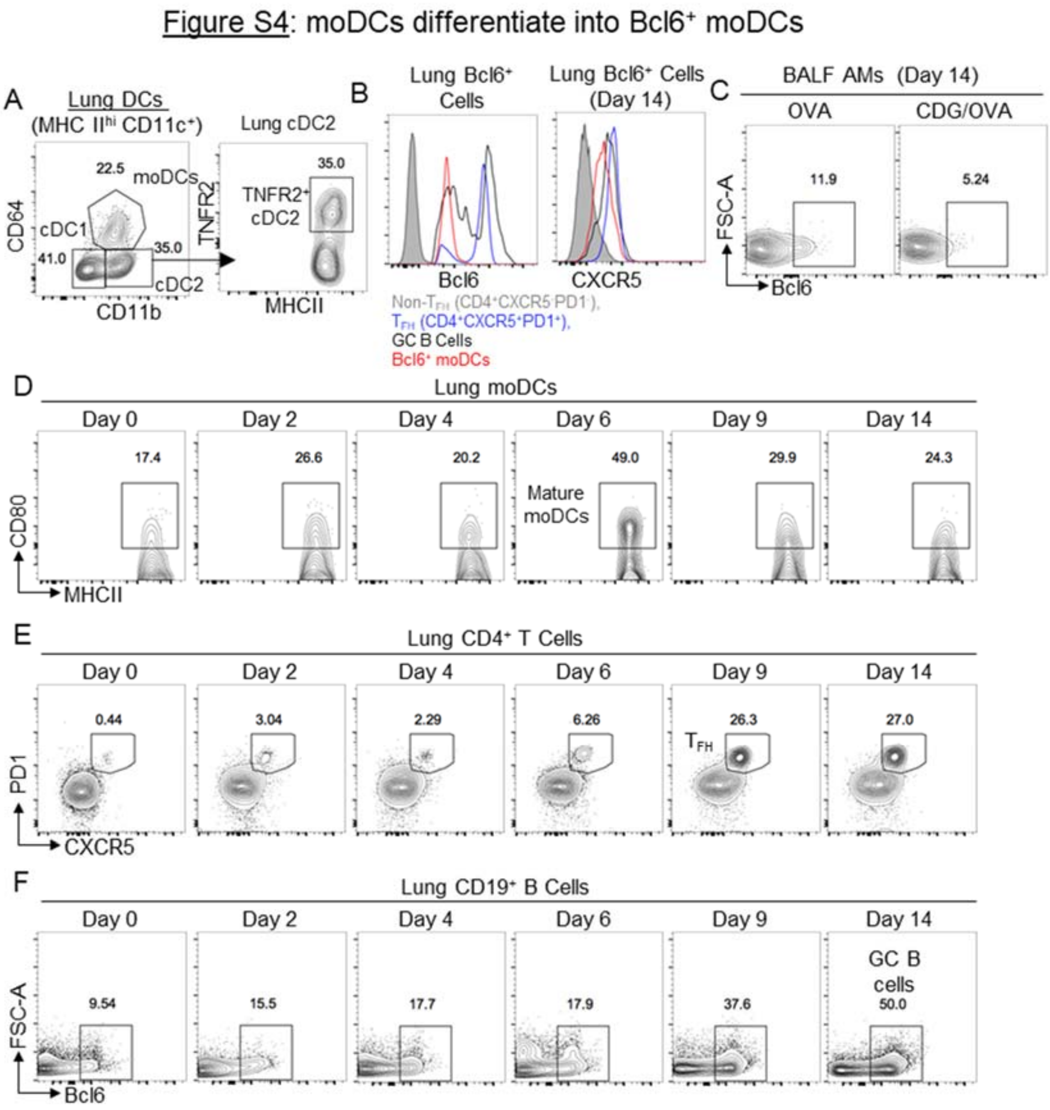


**Figure S4. Lung moDCs differentiate into Bcl6$^+$ moDCs. A.** Gating strategy for lung cDC2. cDC2 are characterized into TNFR2$^+$ and TNFR2$^-$ cDC2 at steady-state. **B.** Bcl6 and CXCR5 expression on indicated cell populations in the lung of C57BL/6J mice on day 14 post CDG/NP$_6$CGG immunization. (n=3mice/group) Data are representative of three independent experiments. **C.** Bcl6 expression in alveolar macrophages in the BALF of WT mice on day 14 post CDG/NP$_6$CGG immunization. (n=3mice/group) Data are representative of three independent experiments. **D-F.** Flow cytometry analysis of CD80$^+$ moDCs (**D**), CD4$^+$PD1$^+$CXCR5$^+$ $T_{FH}$ (**E**), and CD19$^+$Bcl6$^+$ B cells (**F**) of WT mice immunized with CDG/OVA. Lungs were harvested at different time points. (n=3mice/group) Data are representative of three independent experiments.

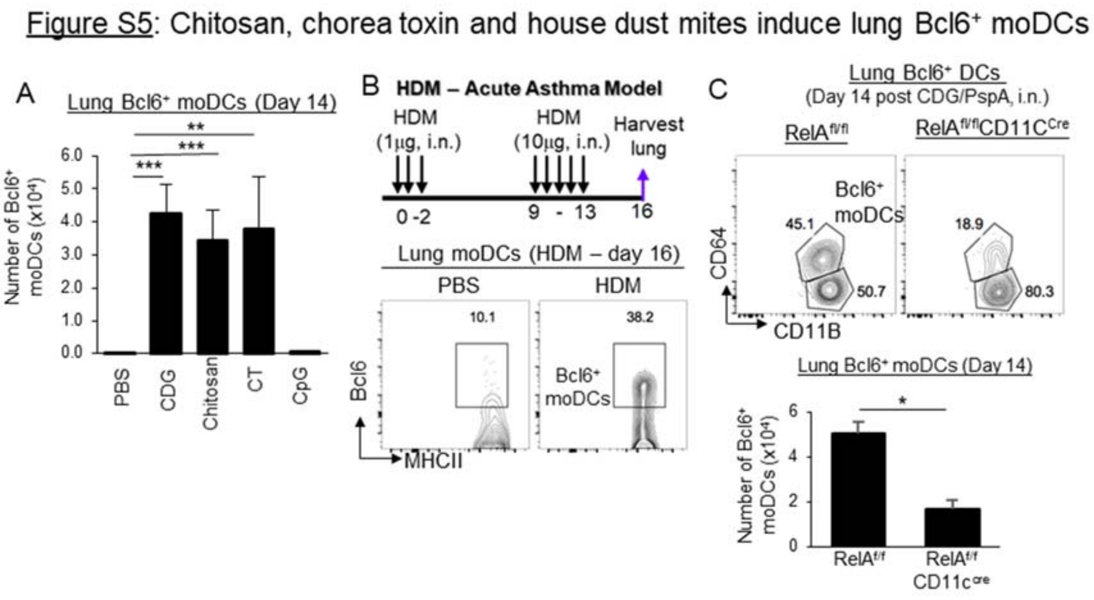


**Figure S5. Chitosan, chorea toxin and house dust mites induce lung Bcl6$^+$ moDCs. A.** WT mice were immunized with NP$_6$CGG (2μg) plus CDG (5μg), chitosan (50μg), cholera toxin (CT) (5μg), and CpG-ODN 2395 (50ng). Lungs were harvested on day 14. Lung Bcl6$^+$ moDCs were enumerated (n=3mice/group) Data are representative of two independent experiments. **B**. Flow cytometry analysis of lung Bcl6$^+$ moDCs in HDM-induced asthmatic WT mice. (n=3mice/group) Data are representative of three independent experiments. **C.** Flow cytometry analysis of lung Bcl6$^+$ DCs in CDG/PspA immunized RelA$^{fl/fl}$ and RelA$^{fl/fl}$CD11c$^{cre}$ mice. (n=3mice/group) Data are representative of three independent experiments. Graphs represent the mean with error bars indication s.e.m. *P* values determined by one-way ANOVA Tukey's multiple comparison test (**A**) or unpaired student *t*-test (**C**). *$P$<0.05, **$P$<0.001, ***$P$<0.0001.

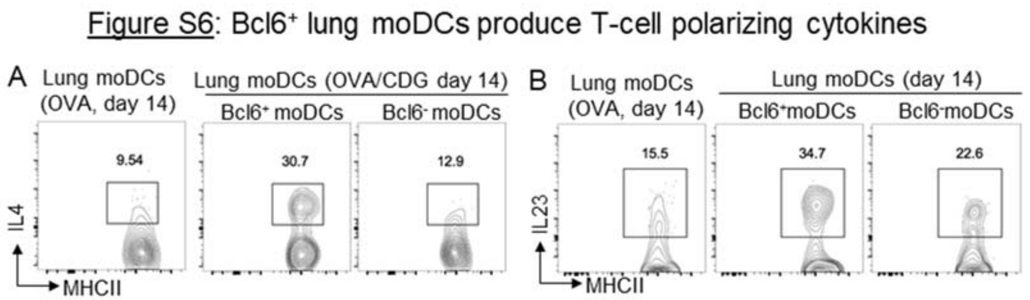


**Figure S6. Bcl6+ lung moDCs produce T-cell polarizing cytokines. A-B.** Flow cytometry analysis of IL-4$^{+}$ (**A**) and IL-23$^{+}$ (**B**) lung moDCs in C57BL/6J mice on day 14 post-immunization (*i.n.*) with CDG/OVA. (n=3mice/group) Data are representative of two independent experiments.

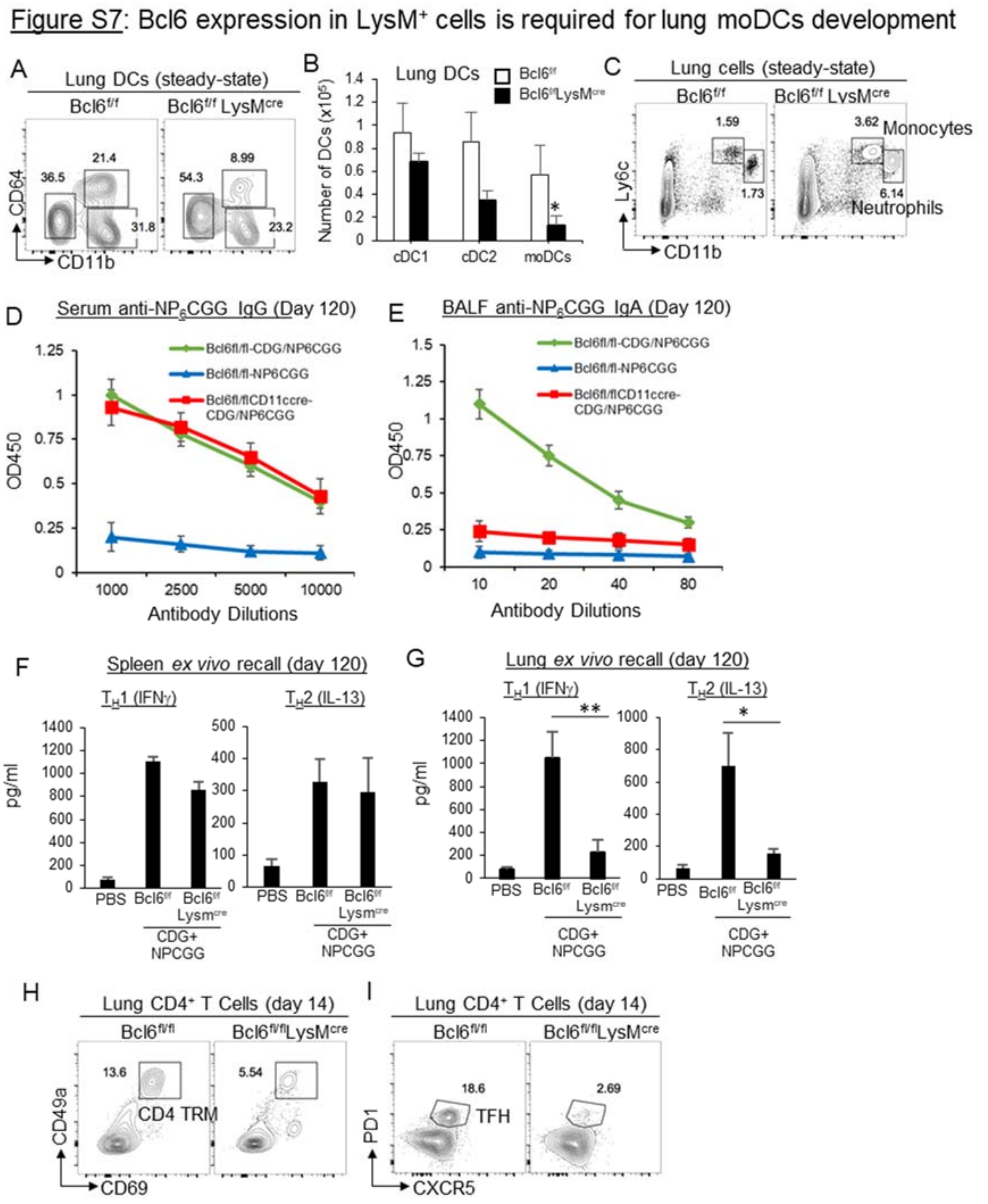


**Figure S7. Bcl6 expression in LysM$^{+}$ cells is required for lung moDCs development. A-B.** Flow cytometry analysis (**A**) and absolute number (**B**) of pulmonary DCs subsets in Bcl6$^{fl/fl}$ and Bcl6$^{fl/fl}$LysM$^{cre}$ mice at steady-state. (n=3 mice/group) Data are representative of two independent experiments. **C.** Flow cytometry analysis lung Ly6C$^{hi}$ monocytes and neutrophils in Bcl6$^{fl/fl}$ and

Bcl6$^{fl/fl}$LysM$^{cre}$ mice at steady-state. (n=3 mice/group) Data are representative of two independent experiments. **D-E.** Bcl6$^{fl/fl}$ and Bcl6$^{fl/fl}$LysM$^{cre}$ mice were immunized with CDG/NP$_6$CGG. Anti-NP$_6$CGG IgG in serum (**D**) and BALF IgA (**E**) were determined by ELISA on day 120 post-immunization. (n=3 mice/group) Data are representative of three independent experiments. **F-G.** Lung cells from immunized Bcl6$^{fl/fl}$ and Bcl6$^{fl/fl}$LysM$^{cre}$ mice in (**D-E)** were recalled with 5μg/ml NP$_6$CGG for 4 days in culture. Cytokines were measured in the supernatant by ELISA. Data are representative of three independent experiments. **H-I**. Bcl6$^{fl/fl}$ and Bcl6$^{fl/fl}$LysM$^{cre}$ were immunized with CDG/NP$_6$CGG. Lung CD4$^+$ T$_{RM}$ cells (**H**) and lung T$_{FH}$ cells (**I**) were determined by Flow cytometry on day 14 post-immunization. (n=3 mice/group) Data are representative of two independent experiments. Graphs represent means ± standard error. The significance is determined by one-way ANOVA Tukey's multiple comparison test (**G**) or unpaired Student's t test (**B**). *$p<0.05$, **$P<0.001$, ***$P<0.0001$.

Figure S8: Bcl6 expression in CD11c⁺ cells is required for lung cDC1 development

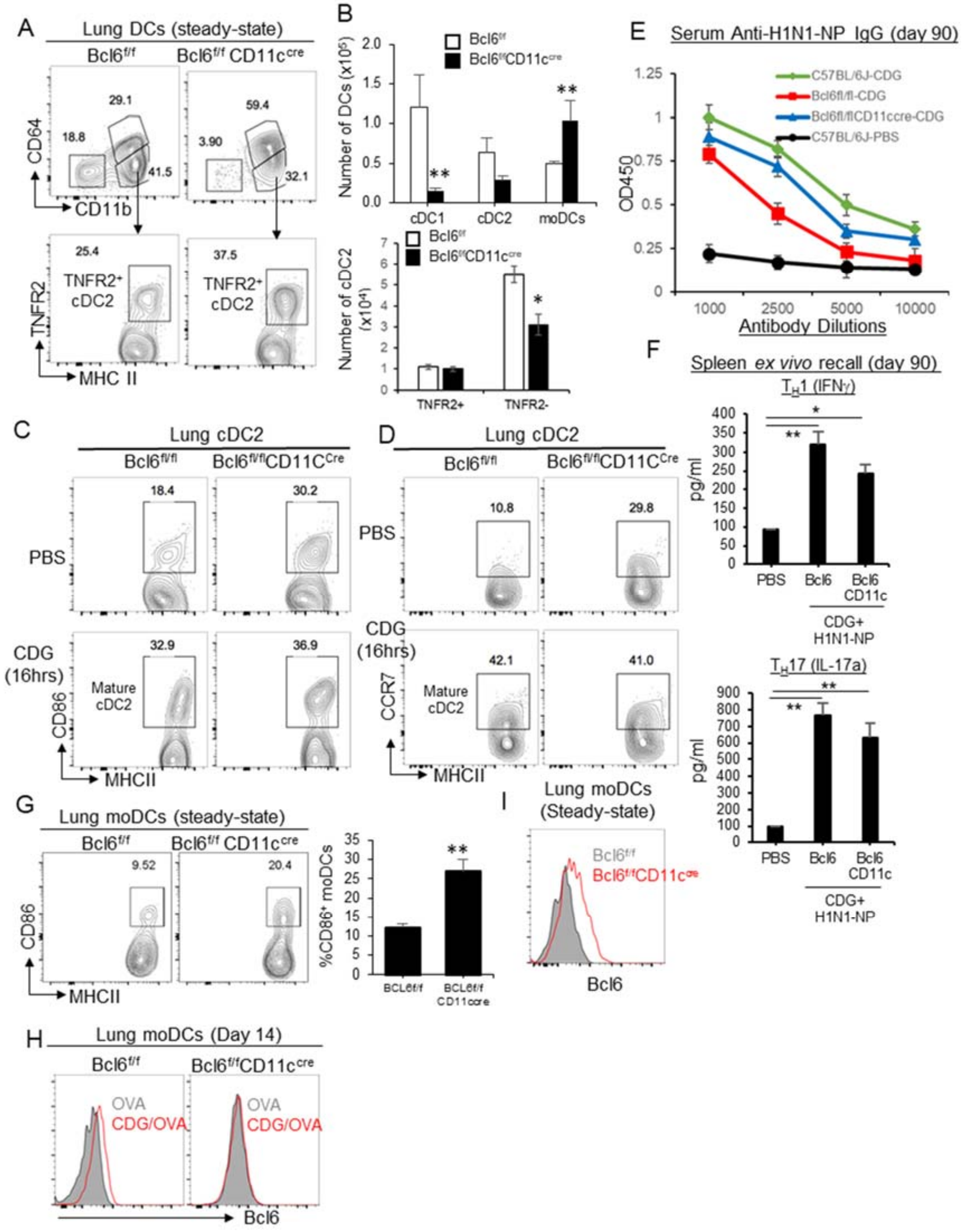


**Figure S8. Bcl6 expression in CD11c⁺ cells is required for lung cDC1 development. A-B.** Flow cytometry analysis (**A**) and absolute number (**B**) of pulmonary DCs subsets in Bcl6$^{fl/fl}$ and Bcl6$^{fl/fl}$CD11c$^{cre}$ mice at steady-state. (n=3mice/group) Data are representative of three

independent experiments. **C-D.** Bcl6$^{fl/fl}$ and Bcl6$^{fl/fl}$CD11c$^{cre}$ mice were treated (i.n.) with CDG (5μg) or PBS for 16hrs. CD86 and CCR7 expression on lung cDC2 were determined by Flow cytometry. (n=3mice/group) Data are representative of two independent experiments. **E.** Bcl6$^{fl/fl}$ and Bcl6$^{fl/fl}$CD11c$^{cre}$ mice were immunized (*i.n.*) with CDG/H1N1-NP twice at two-week interval. Serum anti-H1N1-NP IgG were determined by ELISA on day 90 post-immunization. (n=3mice/group) Data are representative of two independent experiments. **F.** Lung cells from immunized Bcl6$^{fl/fl}$ and Bcl6$^{fl/fl}$CD11c$^{cre}$ mice in (**E)** were recalled with 5μg/ml H1N1-NP for 4 days in culture. Cytokines were measured in the supernatant by ELISA. Data are representative of two independent experiments. **G.** Flow cytometry analysis of CD86 expression on moDCs in Bcl6$^{fl/fl}$ and Bcl6$^{fl/fl}$CD11c$^{cre}$ mice at steady-state. (n=3mice/group) Data are representative of two independent experiments. **H.** Bcl6$^{fl/fl}$ and Bcl6$^{fl/fl}$CD11c$^{cre}$ mice were immunized (*i.n.*) with CDG/OVA or OVA. Bcl6 expression in lung moDCs were determined by Flow cytometry. (n=3mice/group) Data are representative of two independent experiments. **I.** Bcl6 expression on moDCs in Bcl6$^{fl/fl}$ and Bcl6$^{fl/fl}$CD11c$^{cre}$ mice at steady-state. (n=3mice/group) Data are representative of three independent experiments. Graphs represent means ± standard error. The significance is determined by unpaired Student's t test (**B, G**) or one-way ANOVA Tukey's multiple comparison test (**F**). *p<0.05, ***P*<0.001, ****P*<0.0001.

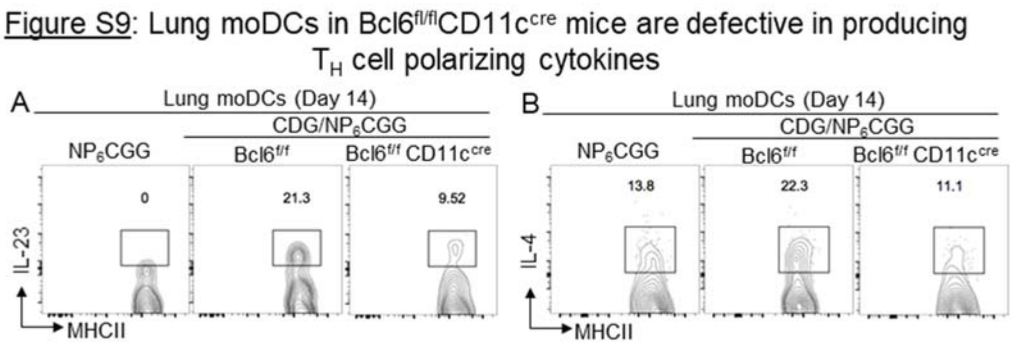


**Figure S9. Lung moDCs in Bcl6$^{fl/fl}$CD11c$^{cre}$ mice lack T$_H$ polarizing cytokines. A-B.** IL-23 (**A**), IL-4 (**B**) production by lung moDCs in Bcl6$^{fl/fl}$ and Bcl6$^{fl/fl}$CD11c$^{cre}$ mice on day 14 post-immunization (*i.n.*) with CDG/NP$_6$CGG. (n=3mice/group) Data are representative of two independent experiments.

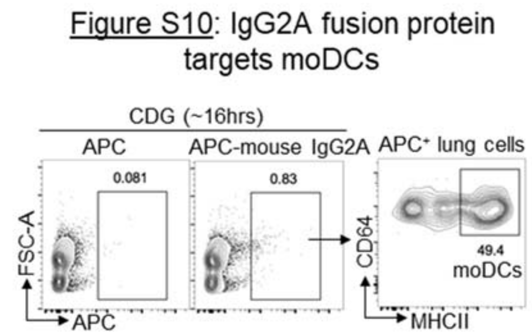


**Figure S10. IgG2A fusion protein targets moDCs.** C57BL/6J mice were administered (*i.n.*) with CDG/APC-mouse IgG2a (clone: MOPC-173) or CDG/APC only. Flow cytometry analysis of $APC^+$ cells were done in the lungs 16 hours post treatment. n=3 mice/group. Data are representative of three independent experiments.
